# Disease-associated NK-cell state remodeling reveals transcriptional profiles predictive of clinical phenotypes

**DOI:** 10.64898/2026.10.22.760478

**Authors:** Fanyi Meng, Peiran Lin, Shunjie Zhang, Tianhao Li, Zili Liu, Peng Ye, Chao Zhong, Yuan Zhou

**Author notes:** Correspondence: Yuan Zhou; Chao Zhong.

## Abstract

Natural killer (NK) cells adapt their cellular states to diverse tissue and inflammatory environments, yet how this plasticity manifests across pathological contexts and relates to patient-level disease phenotypes remains poorly defined. Here we established the Human NK-cell Atlas (HNKA) from over 1 million NK cells, including samples from 1,632 donors spanning 50 diseases. Within a shared NK-cell state architecture, analyses of state abundance and matched-reference exceedance distinguished disease-biased redistribution from transcriptional displacement. Fine-grained analyses localized disease-associated remodeling to specific NK-cell states, including CD56dim NK ISG15 in infection and autoimmunity and CD56bright NK ITM2C in tumors. These patterns were accompanied by recurrent interferon responses and state-specific regulatory activity. Cytokine perturbations recapitulated key transcriptional features of disease-associated NK-cell states. We developed NKceiver, a model based on state-specific NK-cell expression profiles for patient-level prediction. It distinguished 18 diseases and predicted COVID-19 severity and response to immune checkpoint inhibitors, with predictive performance retained in unseen patients and studies. Collectively, our results reveal that disease-associated remodeling within the diverse NK-cell state repertoire resolved by HNKA is biologically interpretable and clinically informative, supporting disease discrimination and prediction of clinical phenotypes.

## Introduction

NK cells are innate lymphocytes that contribute to defense against infection and cancer through cytotoxicity and the production of cytokines and chemokines, while also regulating other immune populations ^1,2^. Their effector activity integrates signals from activating and inhibitory receptors with inflammatory cytokines and the surrounding tissue environment ^3–6^. These responses occur within a heterogeneous repertoire shaped by differentiation, maturation, tissue adaptation and previous immune encounters ^1,7^. This combination of functional specialization and environmental responsiveness makes NK-cell heterogeneity relevant to the balance between protective immunity and immune-mediated pathology, and to differences in disease course and therapeutic response ^2,8^.

Single-cell studies have resolved NK-cell diversity into recurrent populations with distinct differentiation-associated transcriptional programs, transcription-factor activity and functional properties ^1,9^. Tissue adaptation and immune experience contribute additional dimensions, human CD56bright NK cells can acquire tissue-associated features during temporary residence and recirculate through lymphatics, while cytomegalovirus-associated adaptation can involve clonally expanded populations with persistent epigenetic programs ^6,7^. Functional studies in lung cancer further show that tissue-associated NK cells can retain cytotoxic activity and respond to NKG2A blockade ^10^. In SARS-CoV-2 infection, interferon-associated NK-cell activation has been linked to limited antibody breadth; in vitro, interferon stimulation increases NK-cell killing of induced T follicular helper-like cells, connecting inflammatory activation to altered immune regulation ^2^. Analyses of CD8+ T cells have linked state-associated programs to inflammatory and tumor contexts ^11^. It remains unclear how diverse pathological environments shape NK-cell state abundance and transcriptional departures from homeostasis within individual states. Defining which NK-cell states exhibit these transcriptional departures and characterizing the associated regulatory activity could clarify how disease-associated NK-cell remodeling reflects pathological environments and relates to clinical phenotypes.

Because maturation and tissue adaptation generate substantial differences in NK-cell transcriptional programs, the biological meaning of a disease-associated profile depends on the state and tissue against which it is compared ^6,9^. An increase in the representation of a state and a molecular shift in cells occupying that state describe distinct aspects of remodeling. A shared state framework anchored in healthy tissues allows these dimensions to be assessed in relation to normal heterogeneity ^12^. State-resolved transcriptional and regulatory programs can place these departures in the context of NK-cell effector functions and responses to extracellular signals. At the patient level, a shared state framework makes it possible to test whether state-resolved transcriptional profiles capture clinically informative variation, including differences among individuals with the same disease. We therefore sought to define the cellular and molecular organization of disease-associated NK-cell remodeling and establish how state-resolved responses relate to environmental perturbations and patient-level disease phenotypes.

Here we established the HNKA, using the healthy reference of the HNKA to define NK-cell state architecture across tissues. In the disease extension of the HNKA, analysis of state redistribution and matched-reference exceedance revealed that pathological contexts differed in the extent of transcriptional displacement and the fine-grained NK-cell states in which it was concentrated. Recurrent inflammatory programs accompanied the preferential engagement of state-encoded regulatory programs in different disease contexts. Analysis of cytokine-perturbation showed that defined inflammatory stimuli recapitulated NK-cell state transcriptional programs. Building on this state framework, we developed NKceiver to harness patient-level disease information captured by state-resolved transcriptional profiles, enabling prediction of disease identity, COVID-19 severity and response to immune checkpoint inhibitors. These findings support NK-cell states as biologically interpretable cellular readouts that connect pathological remodeling with inflammatory responsiveness and clinically relevant variation.

## Results

### The healthy reference of the HNKA defines cross-tissue NK-cell state architecture

To establish a homeostatic baseline for interpreting NK-cell remodeling across pathological conditions, we assembled a cross-tissue healthy reference of the HNKA from 51 datasets, including 48 published and 3 unpublished datasets. After quality control, the reference comprised 339,615 NK cells from 595 samples spanning 26 tissue types (Fig. 1a,b). We established a three-level annotation hierarchy within each dataset using canonical NK-cell markers, tissue-residency genes and adaptive features. Level 1 captured the canonical CD56dim, CD56bright compartments, level 2 defined major NK-cell classes, and level 3 retained recurrent states reproducible across datasets (Extended Data Fig. 1a,b). Cells and clusters with insufficient marker support or intermediate positions along state continua remained unlabeled.

**Fig. 1.**
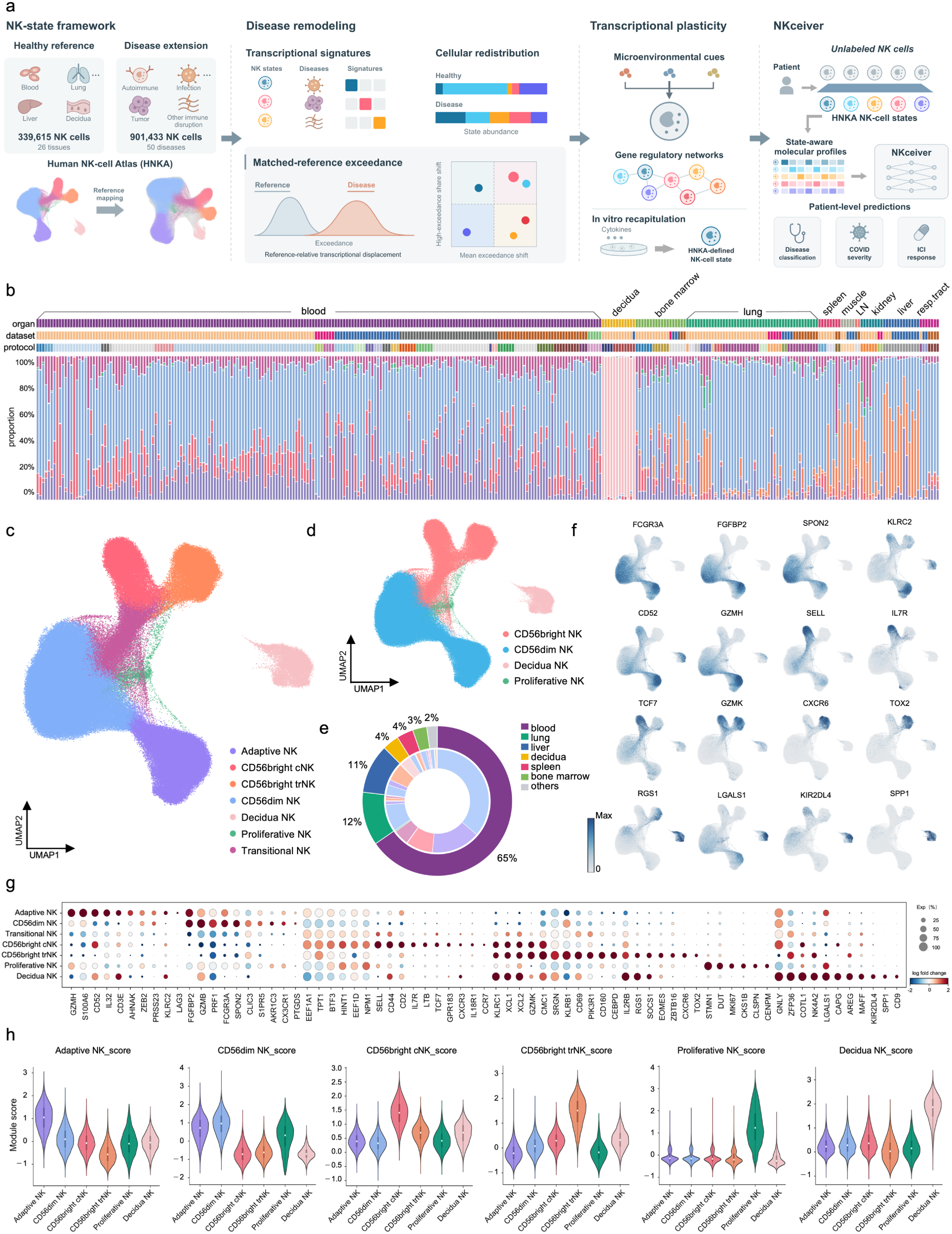
Construction of a cross-tissue human NK-cell reference atlas. **a**, Schematic overview of data collection, atlas construction and downstream applications. **b**, Relative abundance of the major NK-cell compartments in individual samples. Annotation tracks indicate organ, source dataset and sequencing protocol. Each stacked bar represents one sample. **c–d**, UMAP representation of the integrated reference atlas. **e**, Distribution of atlas cells across anatomical sources. The outer ring indicates the proportion contributed by each tissue, and the inner ring shows the corresponding NK-cell composition. **f**, UMAP feature plots showing gene expression of representative markers. **g**, Dot plot of consensus marker genes across the major NK-cell compartments. Dot size indicates the percentage of cells expressing each gene. **h**, Distribution of specific module scores across NK-cell classes. Module scores were calculated from the corresponding consensus marker sets. cNK, conventional NK cells; trNK, tissue-resident NK cells; UMAP, uniform manifold approximation and projection.

To minimize technical variation without obscuring biologically meaningful variations among NK-cell states, we benchmarked six widely used single-cell integration methods using complementary metrics for batch correction and biological conservation ^13–19^ (Extended Data Fig. 1c,d). scPoli with three-level annotation hierarchy provided the best overall performance and enabled label-aware integration, and was therefore used to construct the healthy HNKA embedding. Cells from different studies and sequencing protocols mixed extensively within NK-cell states, with tissue-associated and adaptive populations retaining biologically interpretable local structure (Fig. 1c–e and Extended Data Fig. 1e–g). The integrated reference separated the canonical CD56dim, CD56bright and decidual compartments and resolved seven major classes: CD56dim NK, Adaptive NK, CD56bright cNK, CD56bright trNK, Transitional NK, Decidua NK and Proliferative NK (Fig. 1c,d). Their identities were supported by concordant marker expression and module scores (Fig. 1f–h). CD56bright cNK cells expressed SELL, IL7R, TCF7 and CD44, whereas CD56dim NK cells expressed FCGR3A, CX3CR1, PRF1 and GZMB. Adaptive NK cells combined a CD56dim-like cytotoxic program with KLRC2 and GZMH expression and reduced FCER1G and ZBTB16. CD56bright trNK cells preferentially expressed CXCR6, CD160, TOX2 and CD69, Decidua NK cells expressed CD9, ITGA1 and SPP1, and Proliferative NK cells were marked by MKI67, TOP2A and STMN1 (Fig. 1f,g). Among these major compartments, CD56dim NK, CD56bright cNK and Adaptive NK exhibited marker profiles broadly consistent with the previously described NK1, NK2 and NK3 classification ^1^ (Fig. 1g,h). Beyond these correspondences, the HNKA further distinguished conventional from tissue-associated CD56bright NK cells and resolved Decidua NK as a distinct compartment.

The reference manifold also contained continuous relationships between these classes. Pseudotime analysis identified three major trajectories originating from CD56bright cNK cells (Extended Data Fig. 2a–c), with lineage-associated expression changes distributed along the inferred continuum (Extended Data Fig. 2b). Along the CD56dim trajectory, CCR7 decreased as FGFBP2, S1PR5 and ZEB2 increased, recapitulating progressive acquisition of a mature cytotoxic program (Extended Data Fig. 2d). The CD56bright trNK trajectory was accompanied by coordinated changes in CXCR6, RGS1, EOMES and TOX2, consistent with tissue localization and adaptation (Extended Data Fig. 2e). A third trajectory toward Adaptive NK cells showed progressive loss of FCER1G together with increased KLRC2, ARID5B and PRDM1 (Extended Data Fig. 2f). The healthy HNKA therefore captured recurrent axes of maturation, tissue adaptation and adaptive differentiation within a single reference space for subsequent disease analysis.

### The disease extension of the HNKA reveals shared inflammatory programs and context-dependent state redistribution

We next extended the HNKA across diverse disease environments by curating 901,433 high-quality NK cells from 1,632 donors, 34 major tissue types and 50 diseases (Fig. 2a,b). Diseases were grouped into four broad pathological contexts: autoimmune, infection, tumor and other immune-disruption contexts. The collection included systemic lupus erythematosus (SLE), rheumatoid arthritis (RA), coronavirus disease 2019 (COVID-19), influenza, ovarian cancer (OV), lung adenocarcinoma (LUAD), renal cell carcinoma (RCC), B-cell acute lymphoblastic leukemia (B-ALL), breast cancer (BRCA), colorectal cancer (CRC), chronic obstructive pulmonary disease (COPD), atherosclerosis and other representative immune-mediated conditions.

**Fig. 2.**
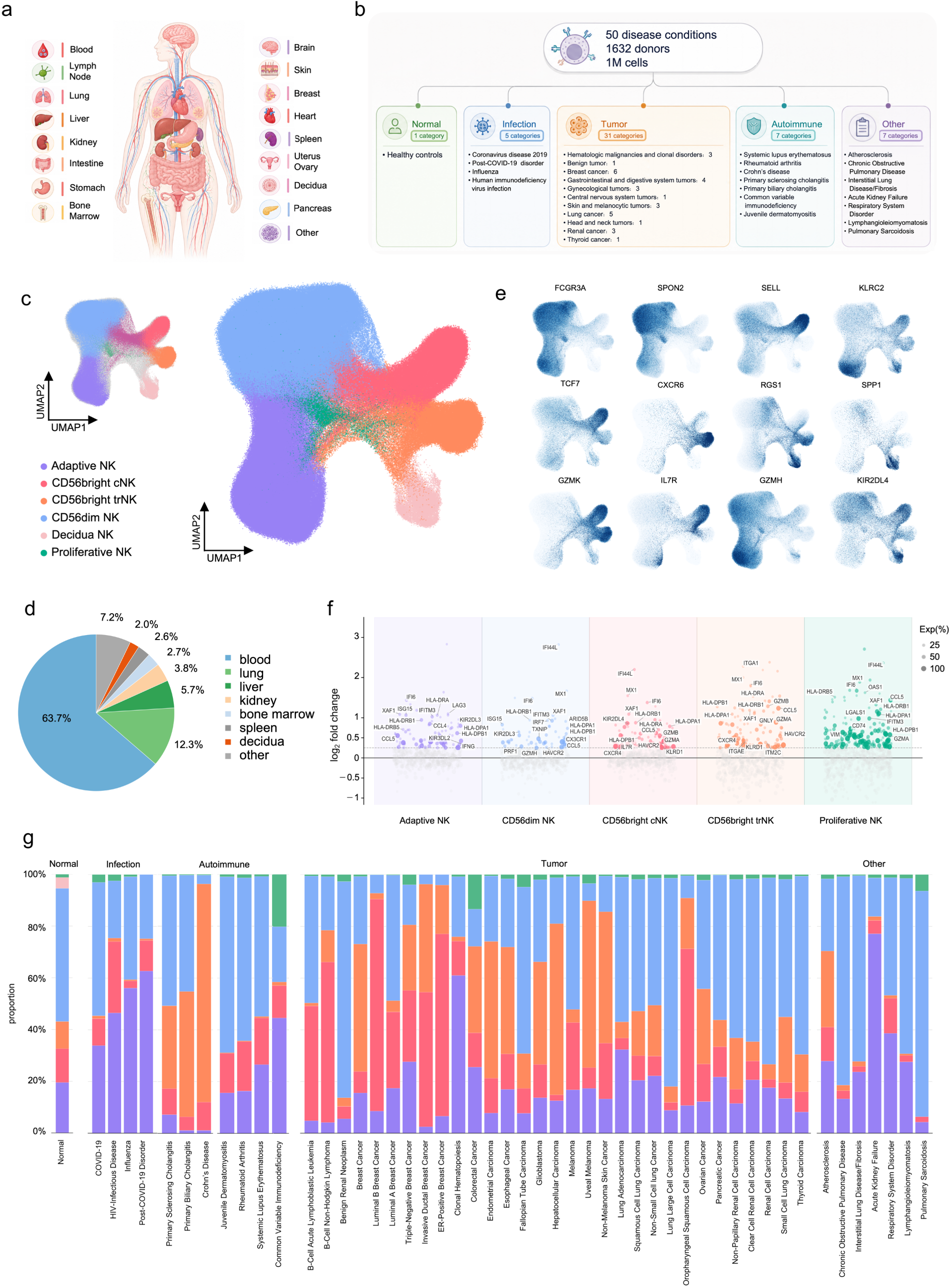
Reference mapping extends the Human NK-cell Atlas across diverse disease contexts. **a**, Anatomical sources represented in the disease extension of the HNKA. **b**, Summary of the disease collection. **c**, UMAP of the HNKA reference colored by major-compartment annotations. **d**, Proportion of mapped cells contributed by the principal anatomical sources. **e**, UMAP feature plots showing expression of representative markers retained after reference mapping. **f**, Differential-expression profiles within the major NK-cell compartments across diseases. **g**, Relative abundance of the major NK-cell compartments across individual diseases, grouped into normal, infection, autoimmune, tumor and other categories.

Disease-derived NK cells were mapped into the healthy HNKA reference while retaining the reference coordinate system. Label transfer assigned cells to six major compartments: CD56dim NK, Adaptive NK, CD56bright cNK, CD56bright trNK, Decidua NK and Proliferative NK, and each retained the core molecular features of its healthy counterpart (Fig. 2c–e and Extended Data Fig. 3a,b). This conservation indicated that major NK-cell identities remained recognizable across disease settings. Compartment-specific marker expression and module-score distributions remained coherent after mapping (Extended Data Fig. 3a,b). Against this stable architecture, tissue- and state-matched comparisons revealed recurrent transcriptional programs superimposed on the mapped identities (Fig. 2f). Interferon-stimulated genes including IFI6, ISG15, IFITM3, XAF1, IRF7 and STAT1 increased across multiple compartments, with strong IFI44L and MX1 induction in CD56dim and CD56bright NK cells. HLA class II genes were induced in parallel, whereas HLA-DRA and CD74 were more selective for Adaptive NK, CD56bright cNK, CD56bright trNK and Proliferative NK cells. These patterns defined a recurrent interferon–HLA class II inflammatory program across distinct NK-cell identities.

Nevertheless, disease contexts exhibited substantial differences in the redistribution patterns of the shared NK-cell repertoire (Fig. 2g and Extended Data Fig. 3c,d). CD56dim NK cells remained the predominant compartment overall, but infection was accompanied by expansion of Adaptive NK cells and reduced representation of CD56bright trNK cells, whereas tumors showed relative enrichment of CD56bright trNK-like cells. In COVID-19, CD56dim NK cells remained predominant while Adaptive NK cells were abundant and increased further in Long COVID. Autoimmune diseases were more heterogeneous, SLE retained a CD56dim-dominant composition, whereas autoimmune cholangitis and inflammatory bowel disease contained more tissue-resident-like compartments. Tumor-associated composition likewise varied with anatomical site and local microenvironment. The summary of major compartments therefore distinguished broad, context-dependent redistribution from organ-associated variation (Extended Data Fig. 3c). Thus, disease-derived NK cells remained organized within the shared HNKA architecture while combining a recurrent inflammatory program with disease-associated remodeling of cellular states.

### Matched-reference exceedance quantifies NK-cell state transcriptional displacement across diseases

Reference mapping provided a common view of NK-cell composition across diseases, but changes in state abundance alone could not determine whether disease-derived cells were also displaced from their corresponding healthy states (Fig. 3a–c). We therefore used matched-reference exceedance to quantify reference-relative transcriptional displacement (Extended Data Fig. 4a). For each mapped disease-derived cell, its distance from the healthy reference was calibrated against the distribution of tissue- and state-matched healthy cells, yielding an exceedance score. At group level, the mean-exceedance shift measured the magnitude of transcriptional displacement, whereas the high-exceedance-share shift identified the compartments contributing disproportionately to the most displaced cells. These two measures allowed disease-associated transcriptional change to be localized without conflating it with state abundance.

**Fig. 3.**
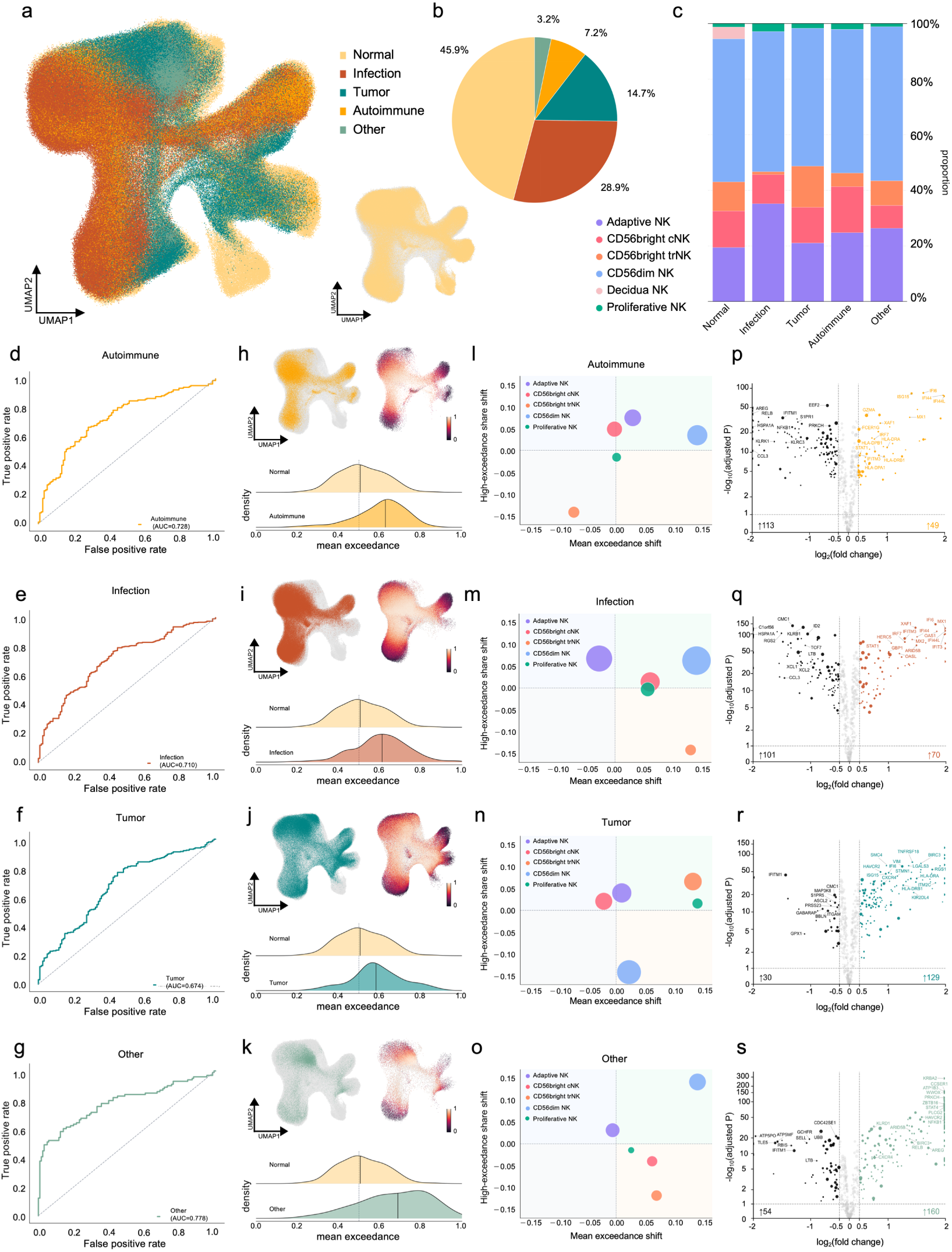
Exceedance resolves disease-associated displacement across NK-cell compartments. **a**, UMAP of the disease extension of the HNKA colored by disease category. The inset highlights healthy reference cells in the embedding. **b**, Relative contributions of normal, infection, tumor, autoimmune and other cells to the mapped atlas. **c**, Relative abundance of the major NK-cell compartments within each disease category. **d–g**, ROC plot evaluating mean exceedance for distinguishing autoimmune (d), infection (e), tumor (f), and other (g) samples from healthy samples. Higher area under the curve (AUC) indicates stronger separation from the healthy reference. **h–k**, Distribution of exceedance in autoimmune (h), infection (i), tumor (j) and other (k) samples. For each category, the left UMAP highlights cells from the indicated category, the right UMAP shows exceedance and the density plots compare mean exceedance with the matched distribution. Vertical lines indicate distribution medians. **l–o**, Major-compartment decomposition of exceedance in autoimmune (l), infection (m), tumor (n) and other (o) disease categories. Each point represents one NK-cell compartment. The x axis shows the mean exceedance shift, defined as the disease–healthy difference in mean atlas exceedance. The y axis shows the high-exceedance share shift, defined as the disease–healthy difference in the fraction of the total high-exceedance cell pool contributed by each compartment. Point size denotes the number of disease-derived cells and color indicates NK-cell identity. Subsets in the upper-right quadrant therefore show both increased mean displacement from the healthy reference and increased contribution to the high-exceedance pool. **p–s**, Differential expression between disease and healthy NK cells in autoimmune (p), infection (q), tumor (r) and other (s) samples.

Mean exceedance distinguished autoimmune, infection, tumor and other disease samples from matched healthy samples by receiver operating characteristic analysis (Fig. 3d–g), and the magnitude and cellular distribution of displacement differed among contexts (Fig. 3h–o). Autoimmune samples showed pronounced displacement, with the largest contributions from CD56dim and Adaptive NK cells (Fig. 3h,l). Autoimmune-associated differentially expressed genes were dominated by inflammatory-response programs (Fig. 3p), and SLE reproduced the category-level pattern in donor-level density, discrimination and compartment decomposition (Extended Data Fig. 4b). Infection-associated displacement was more narrowly concentrated in CD56dim NK cells, with a smaller contribution from CD56bright cNK cells (Fig. 3i,m). The corresponding transcriptional program was dominated by antiviral-response genes, including IFI6, IFI44L, MX1, XAF1, OAS1, IFIT3 and IRF7 (Fig. 3q), and COVID-19 showed the same overall structure (Extended Data Fig. 4c).

Tumors showed lower overall displacement than autoimmune or infection samples, with displacement concentrated in selected compartments (Fig. 3j,n). CD56bright trNK cells made the largest positive contribution, followed by Adaptive NK cells. Proliferative NK cells also showed high mean displacement. The dominant CD56dim compartment remained comparatively close to its matched healthy reference. Tumor-derived NK cells upregulated a distinct set of genes including ITM2C, CXCR4, TNFRSF18, BIRC3, LGALS3, HLA-DRA, RGS1 and KIR2DL4 (Fig. 3r). Ovarian cancer reproduced this CD56bright trNK-dominated pattern at the individual-disease level (Extended Data Fig. 4d). In the other disease category, COPD showed displacement concentrated in CD56dim NK cells (Fig. 3k,o and Extended Data Fig. 4e), accompanied by increased HAVCR2, CXCR4, PLCG2, STAT4 and NFKB1 (Fig. 3s). State redistribution and transcriptional displacement therefore captured separate dimensions of disease-associated NK-cell remodeling. Different pathological environments varied in the magnitude of departure from matched homeostasis, the cellular compartments carrying that departure and the transcriptional programs associated with it.

### Fine-grained NK-cell states localize disease-associated remodeling

To determine whether disease-associated remodeling was broadly distributed within major NK-cell compartments or concentrated in specific subpopulations, we resolved the disease extension of the HNKA into 13 fine-grained NK-cell states (Fig. 4a–d). Within the CD56dim branch, CD56dim NK ISG15 expressed ISG15, IFI6, IFIT3, MX1, OAS1, RSAD2 and STAT1. CD56dim NK CX3CR1 showed CX3CR1, FGFBP2, PRF1, NKG7 and S1PR5, distinguishing interferon-responsive and mature cytotoxic programs. CD56dim NK HAVCR2 was marked by HAVCR2, NEAT1 and CEBPB, and CD56dim NK DNAJB1 by DNAJB1, CREM and DUSP1. Adaptive NK CXCR4 expressed CXCR4, TSC22D3, PRDM1 and TNFAIP3. Adaptive NK KLRC2 retained the KLRC2-associated adaptive program. The CD56bright branch separated into CD56bright NK SELL, expressing SELL, TCF7 and IL7R; CD56bright NK CXCR6, expressing CXCR6, KLRB1, CD69 and GZMK; and CD56bright NK ITM2C, coexpressing ITM2C, CCL5, RGS1, ITGA1 and ZFP36L2. These marker-defined identities were supported by orthogonal functional scoring across the fine-grained states (Extended Data Fig. 5a), reinforcing the functional distinctions inferred from marker genes and module scores. PAGA connectivity further linked CD56bright NK ITM2C with Decidua NK CD9, consistent with partial convergence of tissue-adaptation features between these states (Fig. 4b). Together, graph topology and transcriptional profiles established the fine-grained states as a functional coordinate system for localizing disease-associated remodeling.

**Fig. 4.**
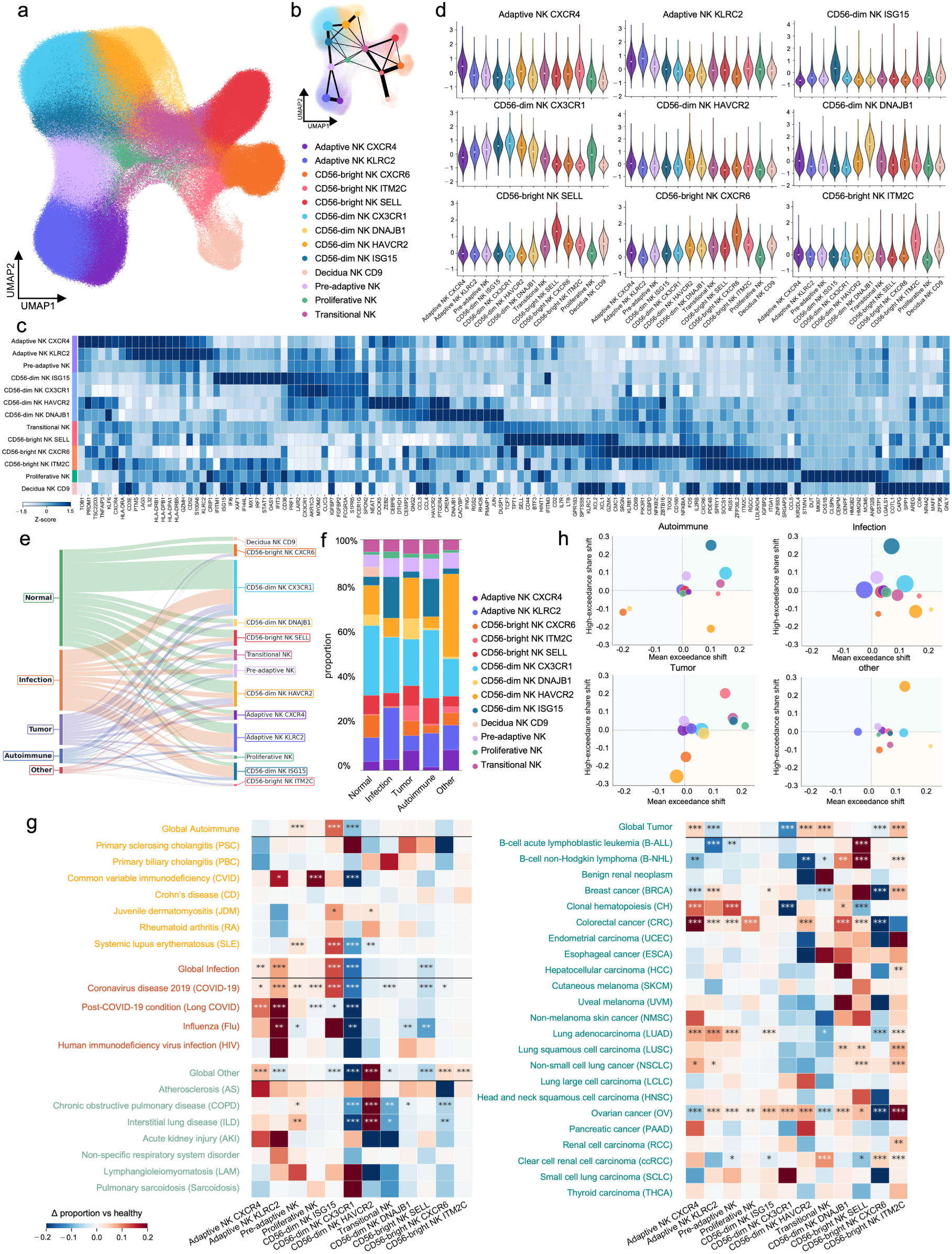
Fine-grained NK-cell states localize disease-associated remodeling. **a**, UMAP of disease extension of the HNKA colored by fine-grained NK-cell states. **b**, PAGA representation of transcriptional connectivity among the NK states. Node size reflects cell abundance and edge width denotes PAGA connectivity. **c**, Heatmap of expression of consensus marker genes across the fine-grained NK-cell states. **d**, Distributions of state-specific marker-module scores across all NK states. **e**, Sankey diagram linking disease categories to fine-grained NK-cell states. Flow width denotes cell abundance. **f**, Relative abundance of NK states within the normal, infection, autoimmune, tumor and other categories. **g**, Tissue-matched changes in NK state proportions between disease and healthy samples. Rows denote global categories or individual diseases and columns denote NK-cell states. (*q < 0.05, **q < 0.01 and ***q < 0.001). **h**, Fine-grained NK-cell states decomposition of exceedance in autoimmune, infection, tumor and other categories. Each point represents one state; the x axis shows the shift in mean exceedance and the y axis shows the change in representation among high-exceedance cells relative to matched healthy samples. Point area denotes relative state abundance.

Fine-grained analysis revealed disease-associated differences in NK-cell state composition within the shared major compartments, with CD56dim NK ISG15 preferentially represented in autoimmune and infection samples and CD56bright NK ITM2C in tumor samples (Fig. 4e,f). Comparisons between diseased and healthy tissues further resolved these patterns (Fig. 4g). CD56dim NK ISG15 increased across autoimmune and infection contexts, including SLE and COVID-19, and CD56dim NK CX3CR1 decreased across multiple diseases. Infection was also accompanied by broad enrichment of Adaptive NK KLRC2. In tumors, CD56bright NK ITM2C was the most consistently increased fine-grained state and was particularly prominent in OV. CD56bright NK CXCR6 decreased across these comparisons. COPD showed a distinct redistribution, with expansion of CD56dim NK HAVCR2 and reduction of the CX3CR1-associated state.

Combining abundance with exceedance further distinguished states with compositional redistribution from those showing concurrent transcriptional displacement (Fig. 4h). In infection, CD56dim NK ISG15 combined increased abundance with a large mean-exceedance shift and a major contribution to the high-exceedance pool; COVID-19 reproduced this coupled pattern (Extended Data Fig. 4c). Autoimmune disease showed a broader configuration in which CD56dim NK ISG15, Adaptive NK and CD56dim NK CX3CR1 contributed to high exceedance (Extended Data Fig. 4b). In tumors, CD56bright NK ITM2C made the largest contribution, with additional displacement in Proliferative and Transitional NK cells, and OV showed the same tumor-associated localization (Extended Data Fig. 4d). Previous studies have shown that the tumor microenvironment can reshape NK-cell transcriptional and functional states ^8,9,20^. An expression signature of CD56bright NK ITM2C was associated with a more favorable progression-free interval in multiple TCGA cancer cohorts (Extended Data Fig. 5b). These observations further linked the tumor-biased ITM2C state to clinically relevant variation across multiple cancer types. In COPD, displacement was concentrated in CD56dim NK HAVCR2 (Extended Data Fig. 4e), consistent with checkpoint-associated remodeling under chronic tissue injury and with sustained NK-cell activation driven by dendritic-cell interactions and IL-15 trans-presentation in the inflamed lung ^21–24^. These results suggest fine-grained HNKA states as the functional units that localized disease-associated compositional redistribution and transcriptional remodeling.

### State-specific regulatory networks exhibit disease-biased activity

We next characterized the regulatory architecture of the fine-grained NK-cell states and its association with disease contexts. Regulon analysis across metacells integrated TF expression, regulon activity and regulon specificity (Fig. 5a). Within the CD56bright branch, CD56bright NK SELL was characterized by a TCF7–LEF1–GATA3 program, CD56bright NK CXCR6 by an NK-lineage and tissue-responsive configuration including NFIL3, EOMES, IRF8 and ETS1, and CD56bright NK ITM2C by a distinct RXRA–ETS2– SIX1-associated program. The CD56dim branch separated into a STAT1–STAT2–IRF7–ETV7 interferon program in CD56dim NK ISG15, a TBX21–KLF2-associated mature program in CD56dim NK CX3CR1, a stress-associated ATF3–DDIT3/AP-1 configuration in CD56dim NK DNAJB1 and a RUNX3–PRDM1– FOXO1-associated network in CD56dim NK HAVCR2. Adaptive NK KLRC2 formed a network centered on TBX21, BATF, IRF5, NFATC3, RELA and STAT6, with additional HOXC5 and ETV2 activity. Adaptive NK CXCR4 combined RUNX3, IKZF1 and IRF1 with RELB and BHLHE40, defining a second adaptive regulatory configuration. Decidua NK CD9 showed a tissue-development program involving NR2F2, RXRA and HOX-family regulators and shared developmental and nuclear-receptor features with CD56bright NK ITM2C. This regulatory convergence paralleled the transcriptional connectivity between the two tissue-adapted states, suggesting that related developmental and microenvironment-responsive modules are reused across distinct NK-cell states.

**Fig. 5.**
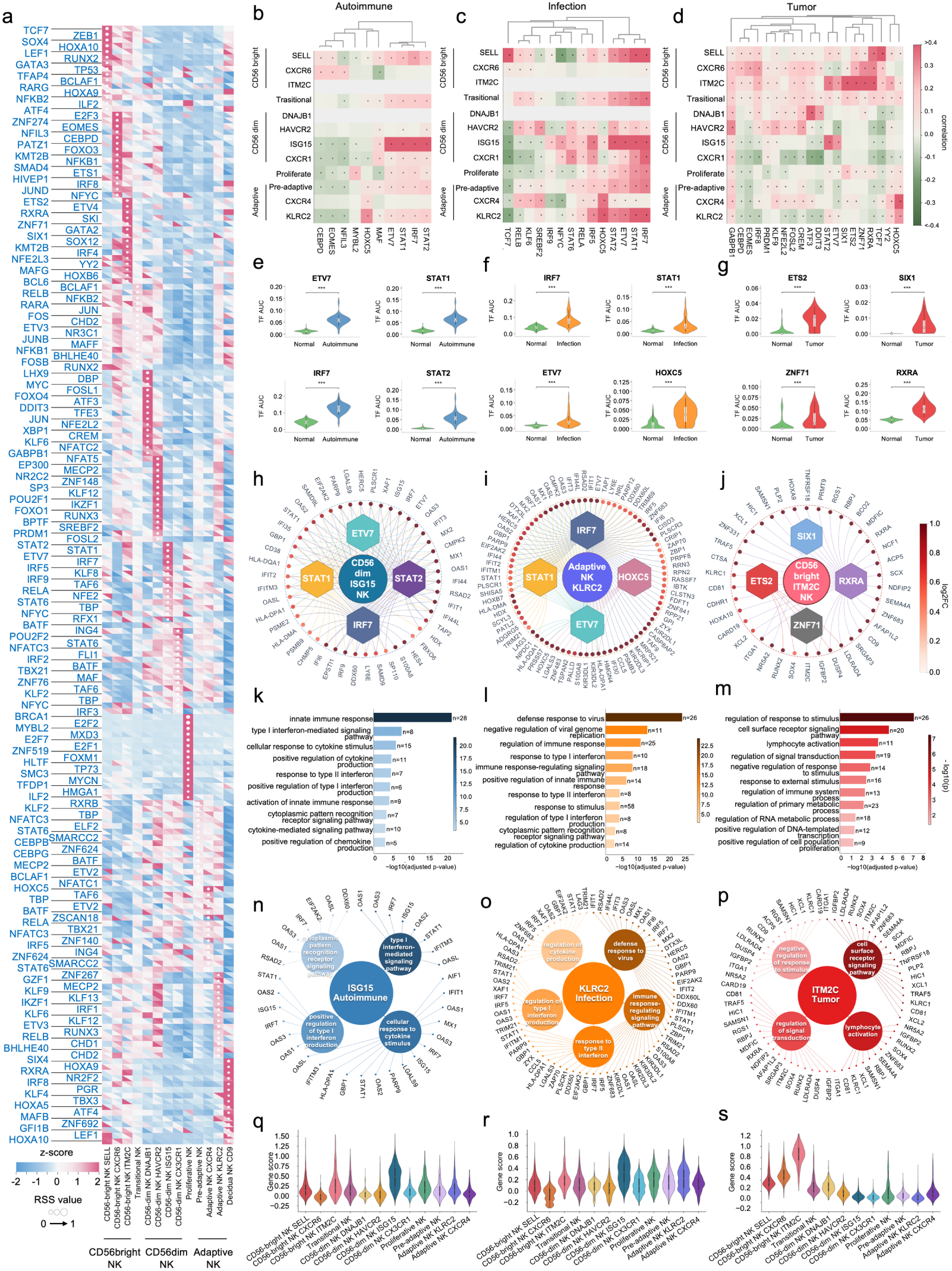
State-specific regulatory networks link NK-cell states to disease-biased remodeling. **a**, Heatmap showing TF expression and regulon activity (AUC) across NK-cell states, with dot size indicating the regulon specificity score (RSS) for the top 15 regulons. Within each tile, the lower-left triangle represents TF expression, whereas the upper-right triangle represents AUC; colors indicate z-scored values. **b–d**, Heatmaps showing correlation between TF regulon activity and disease state across NK-cell states in autoimmunity (b), infection (c), and tumors (d). Color indicates the correlation coefficient; dots mark associations with q value < 0.05. **e–g**, Violin plots showing TF with significant AUC differences between healthy situation and diseases including autoimmune (e), infection (f), and tumor (g) in donor-level by using Mann-Whitney U test with Benjamini-Hochberg FDR correction; *FDR < 0.05, **FDR < 0.01, ***FDR < 0.001. **h–j**, disease-related gene regulatory networks of CD56-dim NK ISG15 in autoimmune (h), Adaptive NK KLRC2 in infection (i), and CD56-bright NK ITM2C in tumors (j) were constructed based on transcription factors whose activity is up-regulated in disease relative to the healthy counterpart, and target genes whose expression is up-regulated with relative disease, with color indicating fold change relative to the healthy counterpart. **k–m**, Bar plot of GO enrichment analyses of the inferred target genes from the representative disease-associated GRNs in autoimmunity (k), infection (l), and tumors (m). Bar length and color indicate −log10(adjusted P value), and numbers indicate the number of target genes enriched in each term. **n–p**, Gene network of GO terms linking selected enriched GO BP terms to the inferred target genes contributing to each term for the representative disease-associated NK cell state. Edges indicate gene-GO term associations. **q–s**, Violin plots of gene-set scores derived from the target genes of the TFs in autoimmunity (q), infection (r), and tumors (s).

Disease-associated regulon activity showed marked state specificity (Fig. 5b–g). In autoimmunity, STAT1, STAT2, IRF7 and ETV7 regulons showed their strongest disease associations in CD56dim NK ISG15, and donor-level analysis confirmed increased activity of these modules (Fig. 5b,e). In infection, Adaptive NK KLRC2 showed increased IRF7, STAT1 and ETV7 activity together with HOXC5 (Fig. 5c,f). Tumors preferentially reinforced RXRA, ETS2, SIX1 and ZNF71 activity within CD56bright NK ITM2C (Fig. 5d,g). The disease contexts associated with these fine-grained states therefore also showed increased activity of their characteristic regulatory programs.

TF-target network reconstruction connected these disease-associated regulons to distinct functional outputs (Fig. 5h–p). In autoimmune-associated CD56dim NK ISG15, STAT1, STAT2, IRF7 and ETV7 formed a highly connected network with shared targets enriched for type I interferon signaling, cellular responses to cytokines and cytoplasmic pattern-recognition-receptor signaling (Fig. 5h,k,n). The infection-associated Adaptive NK KLRC2 network combined an IRF7–STAT1–ETV7 antiviral module with KIR-related receptor programs and targets including ZNF683, CCL5 and ZAP70; enrichment highlighted antiviral defense, immune-response signaling and type I and type II interferon-related functions (Fig. 5i,l,o). Tumor-associated CD56bright NK ITM2C showed a distinct architecture in which RXRA- and SIX1-associated networks included ITGA1, ZNF683, CD9, CD81, RGS1 and CXCR3, alongside XCL1 and XCL2. ETS2- and RXRA-associated targets included signaling-adaptation genes such as IGFBP2, DUSP4 and LDLRAD4, and the ZNF71 regulon contained ITM2C. These targets were enriched for cell-surface receptor signaling, lymphocyte activation and regulation of responses to stimuli (Fig. 5j,m,p). Scoring the disease-associated target modules across all fine-grained HNKA states showed that these networks retained state specificity (Fig. 5q–s). Collectively, these analyses showed that the NK-cell state repertoire captured state-specific patterns of regulatory activity across different pathological environments, reflecting both recurrent interferon responses and context-specific NK-cell remodeling.

### Cytokine perturbations recapitulate transcriptional features of disease-associated NK-cell states

To determine whether defined inflammatory cytokines could reconstruct NK-cell state remodeling, we analyzed an independent cytokine-perturbation dataset ^5^, in which NK cells were exposed acutely or chronically to type I, type II or type III interferons, including IFNβ combined with TNF, IL-6 or IFNγ. Perturbation-derived cells were mapped into the HNKA, assigned to fine-grained states and evaluated by exceedance relative to time-matched unstimulated controls (Fig. 6a,b). The resulting landscape separated cytokine conditions and stimulation durations, enabling direct comparison between experimentally induced responses and HNKA-defined states.

**Fig. 6.**
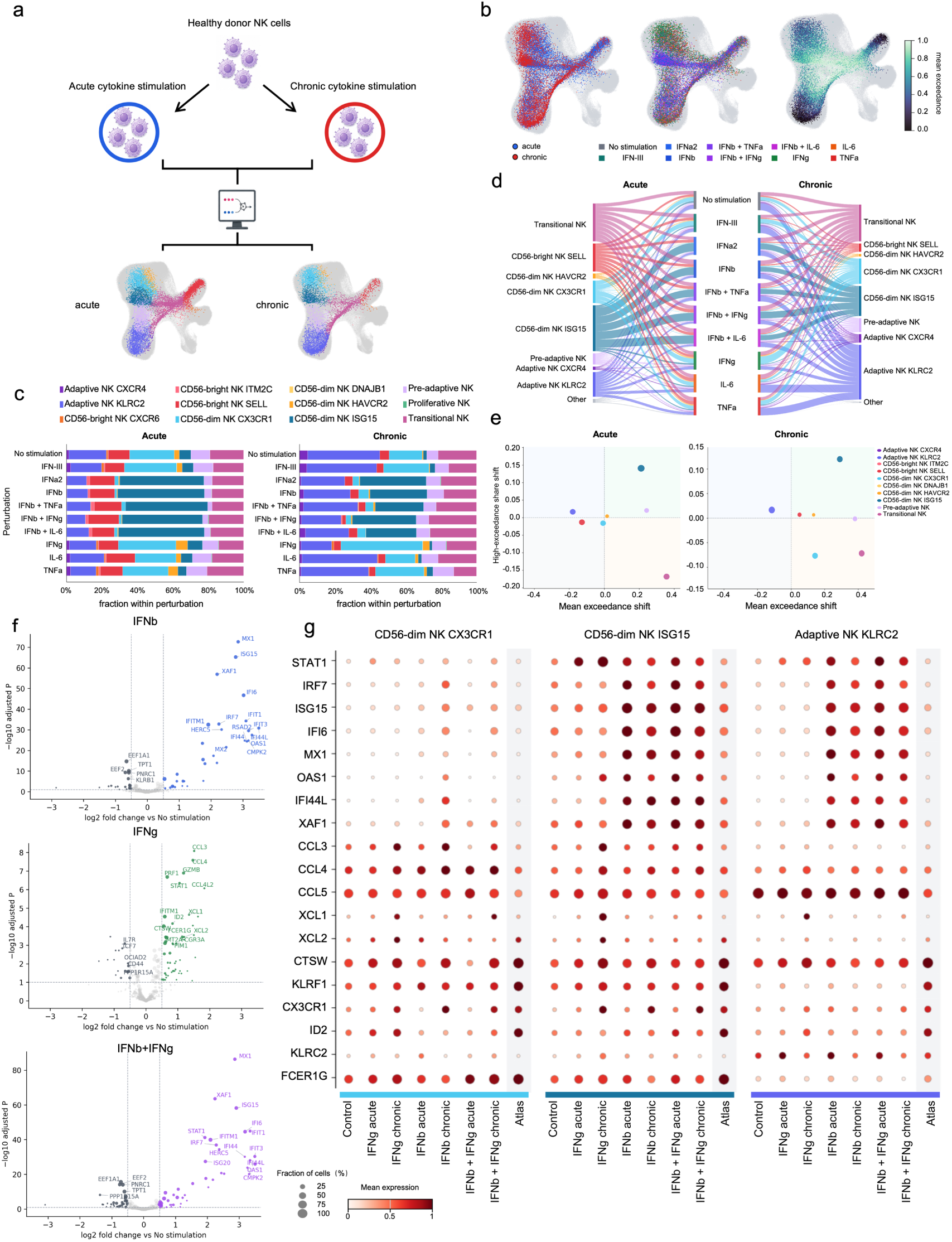
Cytokine perturbations recapitulate transcriptional features of disease-associated NK-cell states. **a**, Experimental and analytical workflow. NK cells from healthy donors were exposed to acute or chronic cytokine stimulation, profiled by single-cell RNA sequencing and mapped onto the disease extension of the HNKA. **b**, UMAP of mapped perturbation-derived cells colored by stimulation duration, cytokine condition and exceedance. **c**, Relative abundance of fine-grained NK-cell states following acute or chronic cytokine stimulation. **d**, Alluvial representation of NK-cell state assignments after acute (left) or chronic (right) cytokine perturbation. Perturbed cells were mapped to the disease extension of the HNKA using soft K-nearest-neighbor assignment. Width denotes the mean mapping proportion from each perturbation condition to the corresponding atlas state; low-abundance assignments are grouped as other states. **e**, Fine-grained NK-cell states decomposition of perturbation-associated displacement after acute or chronic stimulation. The x axis shows the shift in mean exceedance relative to the time-matched unstimulated control, and the y axis shows the corresponding change in high-exceedance representation. Each point represents one state, point area denotes relative state abundance. **f**, Differential expression following IFN-β, IFN-γ or combined IFN-β & IFN-γ stimulation relative to the corresponding time-matched unstimulated controls. **g**, Dot plots comparing the expression of interferon-response, chemokine, cytotoxicity and differentiation-associated genes in representative NK-cell states between the HNKA and cytokine-stimulated samples.

State composition and reference assignment revealed stimulus- and time-dependent patterns of NK-cell states (Fig. 6c,d). Acute IFNα2, IFNβ and IFNβ-containing combinations increased CD56dim NK ISG15 from 2.9% in unstimulated cultures to 43.1–49.3%, with corresponding preferential assignment to this state in the alluvial mapping. IFNγ increased CD56dim NK CX3CR1 from 16.5% to 28.7% after acute exposure and from 20.1% to 45.5% after chronic exposure. Adaptive NK KLRC2 increased across several chronic conditions, type III IFN, IL-6 and TNF cultures, indicating a duration-associated shift.

Exceedance analysis further resolved the cellular distribution of perturbation-associated displacement. Relative to time-matched unstimulated controls, type I interferon conditions produced the largest mean-exceedance shifts, ranging from 0.158–0.185 after acute exposure to 0.208–0.240 after chronic exposure. Chronic IFNγ increased mean exceedance by 0.095, with no clear positive shift after acute exposure, and type III IFN, IL-6 and TNF alone showed smaller changes (Extended Data Fig. 6a,b). State-level decomposition localized these perturbation effects (Fig. 6e and Extended Data Fig. 6e,f). CD56dim NK ISG15 showed the most consistent increase in mean exceedance and contributed disproportionately to the high-exceedance pool. Its displacement increased after chronic type I interferon exposure despite a reduction in relative abundance. CD56dim NK CX3CR1 showed increased displacement and high-exceedance representation under chronic IFNγ.

Differential expression and atlas correspondence analyses further characterized the transcriptional profiles induced by these perturbations (Fig. 6f,g and Extended Data Fig. 6c,d). IFNβ induced an antiviral program involving IFI44L, IFIT1, OAS1, ISG15, IRF7 and XAF1, which was retained under combined IFNβ–IFNγ stimulation with additional STAT1 induction. IFNγ induced a distinct chemokine and mature-effector program marked by CCL3, CCL4, GZMB, PRF1 and CTSW (Fig. 6f). HNKA-state assignment and expression-profile correlation converged on CD56dim NK ISG15 as the closest atlas counterpart to type I interferon-treated cells, supported by coordinated expression of interferon-response genes in mapped cells (Fig. 6g and Extended Data Fig. 6c,d). Together, these perturbation experiments demonstrate that inflammatory cytokine signals can recapitulate transcriptional programs of NK-cell states observed across diverse disease contexts.

### HNKA-defined NK-cell state profiles enable patient-level prediction of disease phenotypes

Having identified disease-selective programs in NK-cell states and shown that defined cytokine stimuli can recapitulate core transcriptional features of disease-associated states, we asked whether these states could provide a readout of patient-level disease phenotypes. We developed NKceiver, a Perceiver-based framework that maps unlabeled NK cells to the HNKA and aggregates their expression into state-specific pseudobulk profiles. Masked cross-attention and a latent Transformer integrate expression from sufficiently supported states for patient-level prediction (Fig. 7a). This representation preserves the shared NK-cell state architecture while retaining patient-specific transcriptional information.

**Fig. 7.**
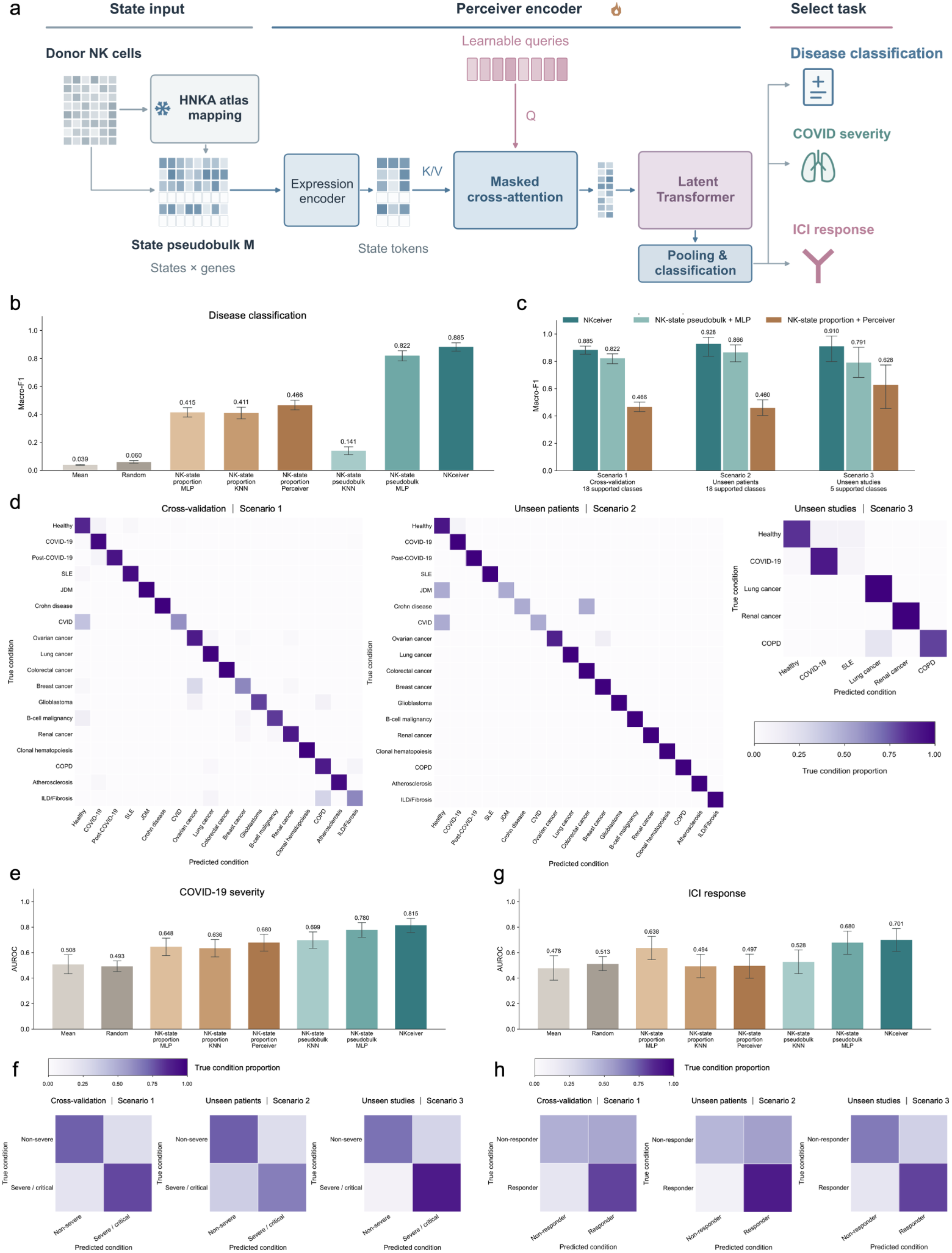
NKceiver predicts disease identity, COVID-19 severity and immunotherapy response from HNKA-defined NK-cell states. **a**, Overview of NKceiver, a Perceiver-based framework for patient-level prediction. Unlabeled NK cells from an individual donor are mapped to the frozen HNKA atlas and aggregated into state-specific pseudobulk profiles. An expression encoder, masked cross-attention and a latent Transformer generate a donor-level representation for task-specific classification. The same architecture is trained separately for each task. **b**, Comparison of NKceiver with alternative input representations and classifiers for disease classification, evaluated by four-fold cross-validation macro-F1. Comparators include NK-state proportions, mean or random expression, and multilayer perceptron (MLP) or k-nearest neighbors (KNN) classifiers. **c**, Macro-F1 of NKceiver, NK-state pseudobulk with MLP and NK-state proportions with Perceiver across cross-validation, unseen patients and unseen studies. Evaluation includes 18 classes in the first two settings and five in unseen studies; predictions retain the full 18-class output. **d**, Confusion matrices of NKceiver for disease classification in cross-validation, unseen patients and unseen studies (left to right; n = 1,215, 242 and 58 patient records, respectively). **e**, Cross-validation area under the receiver operating characteristic curve (AUROC) for COVID-19 severity prediction using NKceiver and the comparator models in b. **f**, Confusion matrices of NKceiver for non-severe versus severe/critical COVID-19, shown in the same order as d (n = 241, 49 and 26 patient records, respectively). **g**, Cross-validation AUROC for immune checkpoint inhibitor (ICI) response prediction using NKceiver and the comparator models in b. **h**, Confusion matrices of NKceiver for ICI responders versus non-responders, shown in the same order as d (n = 150, 30 and 14 patient records, respectively). Error bars indicate 95% confidence intervals from 5,000 patient bootstrap resamples. Confusion matrices are normalized within each true condition.

NKceiver distinguished 18 diseases across 1,215 patient records, achieving a macro-F1 of 0.885 in four-fold cross-validation (Fig. 7b,d). To determine whether this information was explained by redistribution of NK states or also captured their transcriptional features, we compared alternative input representations and classifiers. Using NK-state proportions as input to the same Perceiver architecture yielded a cross-validation macro-F1 of 0.466, compared with 0.885 for state-specific pseudobulk expression (Fig. 7b). Shared mean and random expression inputs also produced lower performance, supporting the contribution of patient-specific transcriptional information. NKceiver ranked highest among the evaluated input and classifier combinations. These comparisons demonstrated that NK-state repertoire captured disease-discriminative information through its transcriptional profiles beyond differences in NK-state abundance.

Disease recognition extended to patients and studies withheld from classifier training. Among 242 unseen patient records spanning all 18 classes, NKceiver achieved a macro-F1 of 0.928 (Fig. 7c,d). Across unseen studies comprising 58 patient records and five conditions, it achieved a macro-F1 of 0.910 while retaining the full 18-class prediction space. NKceiver also outperformed the pseudobulk-based multilayer perceptron and proportion-based Perceiver in both settings (Fig. 7c).

The same state framework further supported prediction of disease severity and treatment response. In COVID-19 and immune checkpoint inhibitor (ICI) response datasets mapped to the HNKA, exceedance analysis identified heterogeneous NK-state displacement between clinical groups (Extended Data Fig. 7a,b). NKceiver distinguished non-severe from severe or critical COVID-19 and ICI responders from non-responders, achieving cross-validation AUROCs of 0.815 and 0.701, respectively, higher than all evaluated comparator approaches (Fig. 7e,g). Discrimination was retained in unseen-patient and unseen-study evaluations, with both clinical classes correctly identified across all three evaluation settings. In the unseen-study evaluation, AU-ROCs were 0.912 for COVID-19 severity and 0.756 for ICI response (Fig. 7f,h and Extended Data Fig. 7c,d). Together, these findings demonstrate that HNKA-defined NK-cell states retain disease-informative transcriptional signatures that can be decoded at the patient level, supporting their potential as cellular biomarkers of disease identity, clinical severity and immunotherapy response.

## Discussion

Our study establishes a shared NK-cell state architecture maintained across diverse pathological environments, with variation in state abundance and transcriptional programs reflecting patient-level disease phenotypes. The HNKA places recurrent inflammatory responses and context-dependent remodeling within this common biological organization. Core features of selected disease-associated programs were recapitulated by cytokine stimulation, and NKceiver read out clinical phenotypes from expression organized by NK-cell state. These observations connect the capacities of NK cells for immune surveillance, tissue adaptation and environmental responsiveness to clinically informative variation.

Previous studies have resolved major human NK-cell subsets, adaptive memory and tissue-associated differentiation, and characterized NK-cell heterogeneity in cancer ^1,8,9^. More recently, a circulating immune-cell atlas linked inflammatory expression profiles to disease classification ^25^. The HNKA integrates maturation, adaptive features, tissue adaptation and inducible functional programs within a fine-grained NK-cell state framework. The healthy reference of the HNKA establishes this biological organization, and the disease extension of the HNKA places diverse pathological responses within it. Matched-reference exceedance complements state redistribution by measuring reference-relative transcriptional displacement against homeostasis. Inferred regulatory programs contextualize this remodeling across the CD56dim, CD56bright and adaptive compartments, revealing inflammatory responses and selective engagement of state-associated programs across diseases. Cytokine stimulation recapitulated the core NK-cell state program observed in disease. The advantage of state-specific expression indicates that molecular variation within NK-cell states carries patient-level disease information. Reliable mapping of unlabeled NK cells to HNKA-defined states enables sufficiently supported expression profiles to be assembled and compared across patients, and predictive value for disease identity, COVID-19 severity and immune checkpoint inhibitor response supports NK-state profiles as candidate cellular biomarkers for several dimensions of clinical phenotype.

Although our study establishes a framework for interpreting the differences and relationships among NK-cell states, several limitations warrant further investigation. The extent to which circulating NK-cell profiles reflect molecular changes within affected tissues remains uncertain. Systemic inflammatory signals and NK-cell trafficking may contribute to concordance between circulating and tissue NK-cell states, whereas local microenvironmental cues may shape organ-specific transcriptional programs that are incompletely reflected in circulation ^6^. This distinction is particularly important for blood-based clinical applications. A further question concerns the mechanisms by which fine-grained NK-cell states might be selectively induced, maintained or redirected. Established examples of NK-cell plasticity provide a basis for investigating these mechanisms ^6,26^, but the signals governing HNKA-defined state changes, the persistence of these changes and their consequences for effector function have yet to be systematically resolved. Distinguishing transient program induction, selective expansion, durable state transitions and changes in cell fate will be essential. Finally, despite patient-level and unseen-study evaluations, the clinical utility of NK-state profiles requires validation in larger, independent, multi-center prospective cohorts across diverse disease populations and treatment settings, particularly when only blood is available.

Translating single-cell profiles into clinical applications requires establishing how faithfully they reflect tissue pathology, track disease activity and predict treatment response across patient populations. The HNKA provides a foundation by organizing disease-associated remodeling within a shared, biologically annotated NK-cell state framework and linking to patient-level phenotypes. Longitudinal profiling of paired blood and tissue samples should clarify when circulating NK-state programs reflect local pathology and how these relationships evolve with disease states. Systematic perturbation of candidate cytokines and regulatory factors, coupled with NK-cell functional assays, could identify the mechanisms governing the induction and maintenance of specific NK-cell states and determine whether transitions between states can be directed or reversed with predictable functional consequences. Ultimately, understanding how NK-cell states reflect pathological environments and how their functions can be deliberately redirected could provide a biological foundation for more precise disease diagnosis and guide targeted strategies for engineering NK cells to enhance immunotherapeutic efficacy.

## Methods

### Dataset collection and metadata harmonization

Human single-cell RNA-sequencing datasets containing NK cells were collected from multiple resources, including the CELLxGENE Portal, the Gene Expression Omnibus (GEO) repository, the Human Cell Atlas (HCA), and unpublished studies (Supplementary Table 1). Datasets were included when an accessible expression matrix and metadata defining tissue and health or disease status were available. Overlapping cohorts were identified from publication, accession, donor, sample and library information and were represented once in the atlas. The healthy reference contained approximately 340K NK cells from 595 samples across 26 tissues. The disease collection initially comprised more than 1 million NK cells, of which 901,433 high-quality cells from 1,632 donors spanning 34 tissues and 50 diseases were retained after quality control and harmonization. Diseases were retained at source resolution and were additionally grouped into autoimmune, infection, tumor and other immune-disruption contexts. Gene identifiers were converted to approved human gene symbols and aligned to GRCh38. Dataset, donor, tissue, sequencing platform, and disease annotations were harmonized to ensure consistent nomenclature and classification across studies.

### Data preprocessing and NK-cell curation

Each dataset was processed independently using study-appropriate quality-control criteria based on library size, detected genes, mitochondrial-transcript fraction and doublet assessment. Raw count matrices were retained for count-based analyses, and log-normalized expression was used for visualization, annotation and signature scoring. Cells with fewer than 200 detected genes, fewer than 500 total counts or more than 20% mitochondrial transcripts were removed. NK cells were selected using source annotations and re-evaluated using canonical NK-lineage markers and cluster-level transcriptional programs. Putative doublets were identified within each dataset using Scrublet v0.2.3 and removed based on doublet scores together with sample-level quality metrics and expression inconsistencies. Clusters exhibiting non-NK cell identities, including T-cell, B-cell, myeloid, erythroid, stromal and epithelial lineages, were removed.

### Hierarchical NK-cell annotation

Initial annotation was performed independently within each healthy dataset using canonical NK-cell markers, cluster-level differential expression and programs associated with cytotoxicity, maturation, adaptive differentiation, tissue residency, interferon response, stress and proliferation. Original study labels were used as supporting information. Cells or clusters with uncertain NK-state identities, including those occupying intermediate transcriptional positions along canonical NK differentiation axes, were assigned as unlabeled. Following integration, annotations were reviewed using marker expression, position in the integrated representation and consistency across donors, tissues and studies. Cluster-associated genes were identified using complementary differential expression analysis implemented in Scanpy, and consensus marker sets were used for cell-state annotation and visualization. NK-cell transcriptional signatures were scored using scanpy.tl.score_genes. Functional programs of NK-cell states were characterized by AUCell scoring of MSigDB C2:CP:WikiPathways gene sets, with state-associated pathways identified by comparison with the remaining NK-cell states. Scores were summarized across NK-cell states, and representative state-associated pathways were selected according to differential activity and biological interpretability. Marker and signature definitions are provided in Supplementary Table 2.

### Integration benchmarking and healthy reference construction

To benchmark and compare single-cell integration methods, PCA, scVI, scANVI, LIGER, Harmony, Scanorama and scPoli ^13–19^ were applied to the reference NK-cell dataset using a common set of 3,000 highly variable genes. Each method was run using the default or officially recommended settings, with method-specific input transformations following the corresponding implementations. Integration performance was evaluated using scIB, which jointly assessed batch correction and conservation of the major NK-cell compartments. scPoli was selected according to its overall scIB performance.

The NK-cell reference atlas was constructed using scPoli with the three hierarchical annotation levels supplied simultaneously as prototype labels. The model was configured with `latent_dim=30`, `embedding_dims=8`, `batch_size=512` and `eta=0.8`. Training was performed for 15 epochs. All remaining parameters followed the default scPoli configuration. The trained model parameters, model gene order, latent representation and hierarchical annotation prototypes were retained as the reference for disease and cytokine-perturbation mapping.

### Topology and differentiation analysis

PAGA ^27^ was used to summarize the connectivity between NK-cell populations. Pseudotime was computed using Palantir ^28^, which captures the continuous nature of differentiation. A cell within the CD56bright NK SELL was selected as the starting cell. Palantir pseudotime was used to compute a cell–cell transition matrix using CellRank ^29^ PseudotimeKernel, and Generalized Perron cluster cluster analysis (GPCCA) was applied to identify macrostates, from which three terminal states were identified. CellRank was subsequently used to estimate each cell’s fate probabilities towards these terminal states. Lineage-associated genes were ranked according to the correlation between their expression and the corresponding terminal-state fate probability. Generalized additive models fitted to cells ordered by Palantir pseudotime were used to calculate gene-expression trends, and lineage-associated genes were displayed in the heatmaps.

### Reference mapping of disease-derived NK cells

Disease datasets were mapped to the established reference using the query-mapping implementation of scPoli. Query expression matrices were aligned to the reference gene order, with genes absent from an individual query represented by zero expression. The trained reference model and annotation prototypes were retained during mapping.

The major NK-cell compartments and fine-grained states were assigned according to the majority label among the 30 nearest reference cells in the scPoli latent space. The latent representation was used for state annotation, exceedance and subsequent comparisons of NK-cell composition and transcriptional programs across disease conditions.

### Compositional analysis

Cell-state abundance was calculated within donor–tissue–condition units. Multiple samples from the same donor, tissue and disease condition were combined before calculating abundance. Disease-versus-healthy comparisons required at least five donors and 1000 NK cells in each group. Disease and healthy units were compared within matched tissues or organs using harmonized anatomical annotations. Abundance changes were reported as percentage-point differences between mean donor-level proportions and tested using two-sided Mann–Whitney U-tests. P values were adjusted using the Benjamini–Hochberg procedure separately within each disease context and annotation resolution. Broad disease-context comparisons and individual-diagnosis comparisons were treated as separate analysis families. Sankey diagrams were generated from mapped cell counts as descriptive summaries, whereas compositional heatmaps show tissue-matched differences in donor-level abundance.

### Exceedance framework

Exceedance quantified each NK cell’s displacement from the source atlas relative to a matched control distribution. In the fixed latent space, the reference distance was the mean Euclidean distance to its *k* nearest source-atlas cells:

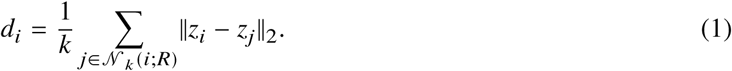

Here, *z_i_* denotes the latent representation of cell *i*, *R* the source-atlas reference and *N_k_*(*i*; *R*) its nearest neighbors. Distances were converted to empirical control-relative percentiles:

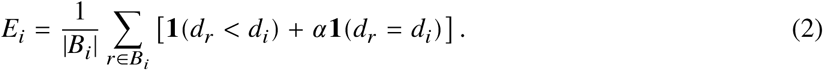

Here, *B_i_* contains the calibration control cells. Higher scores indicate greater reference-relative displacement. For disease analyses, normal controls were matched by major NK-cell compartment and organ, requiring at least 100 cells from five donors. For perturbations, mapped cells with annotation support below 0.60 were excluded before calibration. No-stimulation controls were then pooled within each time point, requiring at least 20 cells from two donors.

Mean exceedance was calculated within donor–condition units containing at least 20 eligible cells. Disease density and receiver-operating-characteristic analyses used these unit means. Perturbation densities instead used cell-level scores, with equal total weight per donor, a Gaussian bandwidth of 0.05 and peak-normalized curves; displayed means were averages of donor-specific means. Overall perturbation shifts were calculated as the mean within-donor difference from time-matched No-stimulation controls among paired donors.

For disease state maps, the mean exceedance shift was the difference between disease and normal pooled-cell means within the same compartment or state. Donor-level statistical comparisons required at least ten cells per state unit and ten eligible units per group. For perturbation state maps, state-specific means were averaged equally across eligible non-control perturbation units and centered on the timepoint-wide mean of No-stimulation donor means across all states. States required at least 50 non-control cells from two donors, four eligible units containing at least five state cells each, and median annotation support of at least 0.75.

High-exceedance share measured the fraction of the high-exceedance pool assigned to each state:

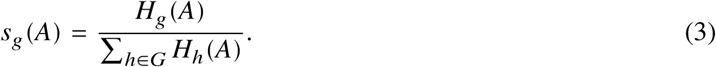

Here, *H_g_*(*A*) is the number of high-exceedance cells in state *g* within group or unit *A*, and *G* is the set of states contributing to the denominator. In the disease atlas, high-exceedance cells were defined as those with scores above the 90th percentile of the normal-control exceedance distribution in each comparison. The high-exceedance share shift was the difference between disease and normal groups in the fraction of this pool assigned to each state. For perturbations, high-exceedance cells were defined as those with scores of at least 0.90. Shares were calculated within each donor–perturbation unit over the eligible state set. Shares were then averaged equally across non-control units and, separately, No-stimulation donor units. The non-control mean minus the control mean defined the high-exceedance share shift. State maps displayed mean exceedance shift on the x-axis and high-exceedance share shift on the y-axis.

### Survival analysis

The prognostic relevance of the CD56bright ITM2C state was assessed in The Cancer Genome Atlas (TCGA) cohorts using GEPIA3^30^. An ITM2C-state signature comprising ITM2C, CD9, ITGA1, KIR2DL4 and SOCS1 was derived from the marker genes used to define this state in the HNKA. For each cancer type, a signature score was calculated using the first principal component of the expression profiles of these genes. Patients in the upper and lower quartiles of signature scores were assigned to high- and low-signature groups, respectively. Progression-free interval (PFI) was evaluated using Kaplan–Meier analysis, with differences between groups assessed by the log-rank test. Hazard ratios (HRs) and 95% confidence intervals were estimated using Cox proportional-hazards analysis. Analyses were performed separately for each TCGA cancer type.

### Metacell construction

Metacells were constructed using SEACells ^31^ independently for each NK cell state per sample. The target compression ratio was approximately 50 cells per metacell, assigning SEACell_IDs. For NK cell states with fewer than 100 cells in a sample, all cells were assigned the same SEACell_ID. SEACell_IDs were stored in the metadata. The “SEACells.core.summarize_by_SEACell” function was used to aggregate raw counts from cells assigned to the same SEACell label. This process generated 22,569 NK metacells.

### Construction of transcription factor regulatory networks

TF-target gene regulatory networks were inferred from scRNA-seq metacell expression profiles using pySCENIC ^32^. Co-expression-based TF-target adjacencies were first inferred using GRNBoost2^33^, with the filtered expression matrix as input and the retained TFs as candidate regulators. Motif enrichment and target pruning were performed using human cisTarget motif-ranking databases for hg38, and high-confidence regulons were subsequently generated using pyscenic.prune.df2regulons. This procedure identified 87 motif-supported regulons, whose activities were quantified across 22,569 metacells using AUCell and stored for downstream regulon activity analyses.

### Downstream analysis of GRNs

To investigate the downstream regulatory programs of NK cell GRNs, regulon specificity scores (RSS) were first visualized as network plots to evaluate the global variation of transcription factor activity across different NK cell states. Disease-associated transcription factor activity was then systematically assessed for each NK cell state. AUCell activity of each regulon was correlated across metacells with disease status by point-biserial Pearson correlation, separately for autoimmune, infectious and tumor conditions. P values were adjusted by the Benjamini–Hochberg procedure within each condition, and the regulons with the strongest positive association per condition (q < 0.05) were retained. From these, biologically relevant TF modules were selected for further analysis. For each TF module, GRN network plots were generated to characterize the regulatory architecture of the corresponding NK cell under disease conditions, including the interactions between transcription factors and their predicted downstream target genes. To further quantify disease-associated signatures, gene score analysis was implemented to score each NK cell state based on disease-specific gene sets, enabling a systematic evaluation of specific expression patterns.

To further explore the functional implications of these regulatory programs, target genes regulated by TF modules were subjected to gene ontology enrichment analysis using g:Profiler ^34^, with all annotated genes as the statistical background and multiple-testing correction by the g:SCS method. Enriched GO terms were used to infer the biological pathways and functional roles mediated by disease-associated regulatory networks.

### Perturbation mapping and analysis

NK cells were extracted from the processed cytokine-stimulation dataset ^5^, comprising cells from three healthy donors with 2 or 7 days of stimulation. Donor, treatment and time point annotations provided with the published dataset were retained. Perturbation-response NK cells were mapped to the integrated NK-cell atlas using the pretrained scPoli reference model described above. For each mapped cell, nearest atlas neighbors were identified in the scPoli latent space, and the proportion of neighbors belonging to each fine-grained atlas state was retained as a soft state-association vector. Exceedance was calculated relative to unstimulated NK cells from the corresponding sampling time using the procedure described above.

For state-association analysis, cells were grouped by donor, perturbation, time point and mapped fine-grained state. Soft state-association vectors were averaged within each unit and subsequently averaged with equal weight across units to obtain the atlas-state association profile of each mapped query state. Transcriptional similarity to atlas states was assessed independently of latent-space association. Query and atlas counts were library-size normalized and log transformed over the highly variable genes shared between the datasets. Atlas and query profiles were centered relative to the equal-weighted mean profile across atlas states. For each query unit, Pearson correlations were calculated between its centered expression profile and each atlas-state centroid across the selected genes, and correlations were combined across units after Fisher z transformation. Perturbation-level state distributions were summarized from the cell-level soft association vectors. Acute and chronic perturbations were visualized separately using alluvial diagrams, with flow width proportional to the donor-balanced assignment mass linking each perturbation to each fine-grained atlas state.

### NKceiver: NK-state representation and patient-level prediction

We developed NKceiver, a patient-level classification framework based on NK-state pseudobulk expression, for disease classification, COVID-19 severity classification and ICI response classification. Dataset accessions are summarized in Supplementary Table 3. NK cells were assigned to HNKA states using a fixed panel of 3,000 HVGs. A frozen expression-to-latent mapping network projected normalized cell expression into the HNKA reference space, and state labels were transferred by distance-weighted voting among the 15 nearest reference neighbors. Query samples were processed using the fixed mapping network.

For each original sample *s*, let *x_ig_* denote the raw count of gene *g* in cell *i*, and let *C_sk_* denote the set of cells assigned to NK state *k*. We constructed the state pseudobulk matrix **M***_s_* ∈ ℝ*^K^*^×*G*^, by summing raw counts within each state, normalizing over the fixed gene panel and applying a log1p transformation:

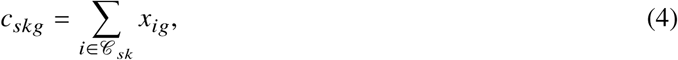

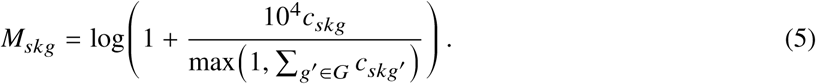

Here, *M_skg_* is the normalized pseudobulk expression of gene *g* in state *k* of sample *s*. Only valid state profiles contributed to classification; samples containing at least one valid state were eligible for prediction. Missing or low-support states were excluded through an attention mask rather than assigned imputed expression profiles.

NKceiver used a compact Perceiver-based architecture ^35^. An expression encoder projected valid NK-state pseudobulk profiles into 128-dimensional tokens. Eight trainable queries aggregated these tokens through four-head cross-attention with separate query and context normalization, a query residual connection and a feed-forward residual block. A single latent Transformer block was followed by mean pooling, a 128-dimensional projection and a task-specific classification head. State validity was enforced through an attention mask.

The same architecture was fitted separately for the three tasks. For each patient *d*, class probabilities were averaged equally across usable original samples:

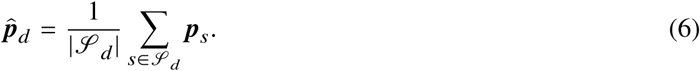

Here, *S_d_* denotes the usable samples from patient *d*, and ***p****_s_* is the class-probability vector for sample *s*. Disease predictions used the highest probability among all 18 categories. COVID-19 severity and ICI response predictions used a fixed threshold of 0.5 for the severe/critical and responder probabilities, respectively.

### Model training and evaluation

Neural classifiers were trained with AdamW for a maximum of 1,500 optimization steps using three fixed random seeds. Sample-level cross-entropy losses were averaged within patients before patient-level aggregation. Disease classification used class weights proportional to the square root of inverse training-class frequency, normalized to unit mean. COVID-19 classification used natural patient sampling without class weighting; ICI classification used label-balanced patient sampling without additional class weights. Model selection and training duration were determined using internal validation within each training pool, followed by refitting on the corresponding training data.

Three evaluation protocols were used. Cross-validation comprised four patient-level folds within the development pool after exclusion of designated studies. Unseen-patient evaluation reserved approximately 20% of this pool, with development selection and refitting restricted to the remaining patients. Unseen-study evaluation trained on the development pool and evaluated jointly withheld studies. Each protocol used separately fitted classifiers, and evaluation records were excluded from classifier fitting, preprocessing estimation and internal model selection. Disease study holdouts comprised GSE161918, PRJNA705464, GSE173896 and phs002371. COVID-19 holdouts comprised EGAD00001007718 and GSE149689. ICI holdouts comprised GSE235863 and GSE206325.

### Comparator models and statistical analysis

Cross-validation compared eight configurations: Mean and Random expression controls; NK-state proportions with a multilayer perceptron (MLP), k-nearest neighbors (KNN) or Perceiver; and NK-state pseudobulk with each of these three classifiers. Both expression controls were processed with the same panel normalization and log1p transformation. State proportions used all eligible NK cells as the denominator, including cells in low-support states. Comparators shared evaluation patients and folds, with training-only parameter selection.

Disease performance was assessed primarily by macro-F1. COVID-19 and ICI performance was assessed primarily by the area under the receiver operating characteristic curve. Additional measures included accuracy, balanced accuracy and class-specific recall. Study-holdout macro-F1 was summarized over the classes represented in the evaluation set, with the class range reported. Confusion matrices were normalized within true classes. Confidence intervals quantified patient-sampling variability conditional on the fitted predictions.

## Acknowledgements

We thank Shanghai Artificial Intelligence Laboratory for its support. This study was supported by the National Natural Science Foundation of China (grant no. 32570789 to Y.Z.), Peking University Medicine plus X Pilot Program-Artificial Intelligence and Medical Development Initiative (grant no. BMU2025YXXLHAIYX022 to Y.Z.).

## Author contributions

Conceptualization: Fanyi Meng, Yuan Zhou; Methodology: Fanyi Meng, Yuan Zhou; Investigation: Fanyi Meng; Data Curation: Fanyi Meng, Peiran Lin, Shunjie Zhang, Tianhao Li; Formal Analysis: Fanyi Meng, Peiran Lin, Shunjie Zhang, Zili Liu, Peng Ye; Supervision: Yuan Zhou, Chao Zhong; Writing - Original Draft: Fanyi Meng; Writing - Review & Editing: Yuan Zhou, Chao Zhong.

---

**Extended Data Fig. 1.**
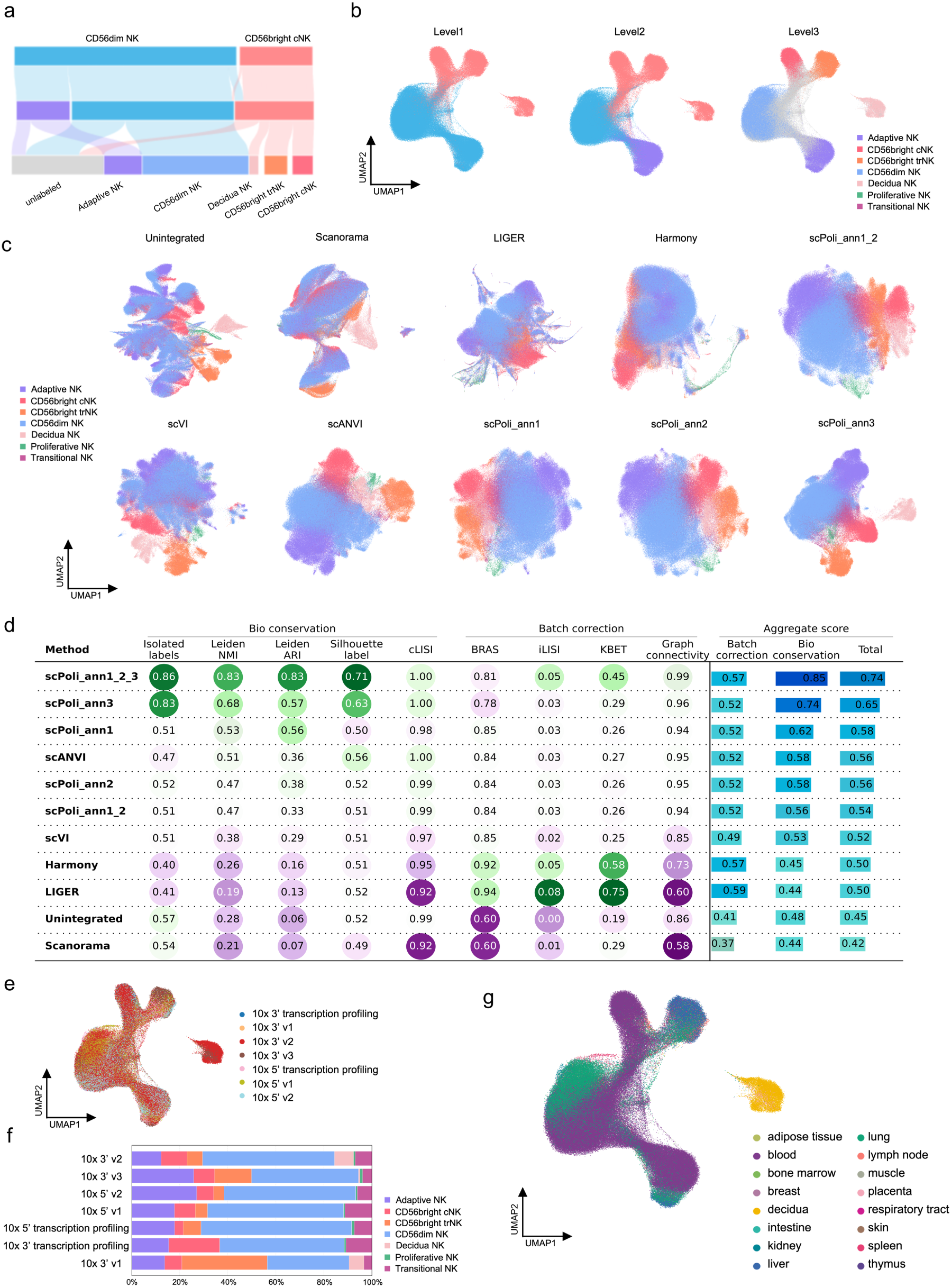
Hierarchical annotation and integration benchmarking of the reference NK-cell atlas. **a**, Sankey diagram linking hierarchical annotations used for label-aware integration. Flow width denotes cell abundance. **b**, UMAP of the HNKA colored according to the level 1, level 2 and level 3 hierarchical annotations used for label-aware integration. **c**, UMAP comparison of unintegrated data and representations generated using Scanorama, LIGER, Harmony, scVI, scANVI and alternative scPoli annotation configurations. **d**, Quantitative comparison of integration methods using complementary biological-conservation and batch-correction metrics. Circle size and color denote normalized metric values. Aggregate biological-conservation, batch-correction and total scores are shown on the right. **e**, UMAP of the representation colored by sequencing protocol. **f**, Relative abundance of the major NK-cell compartments within each sequencing protocol. **g**, UMAP of the representation colored by tissue source.

**Extended Data Fig. 2.**
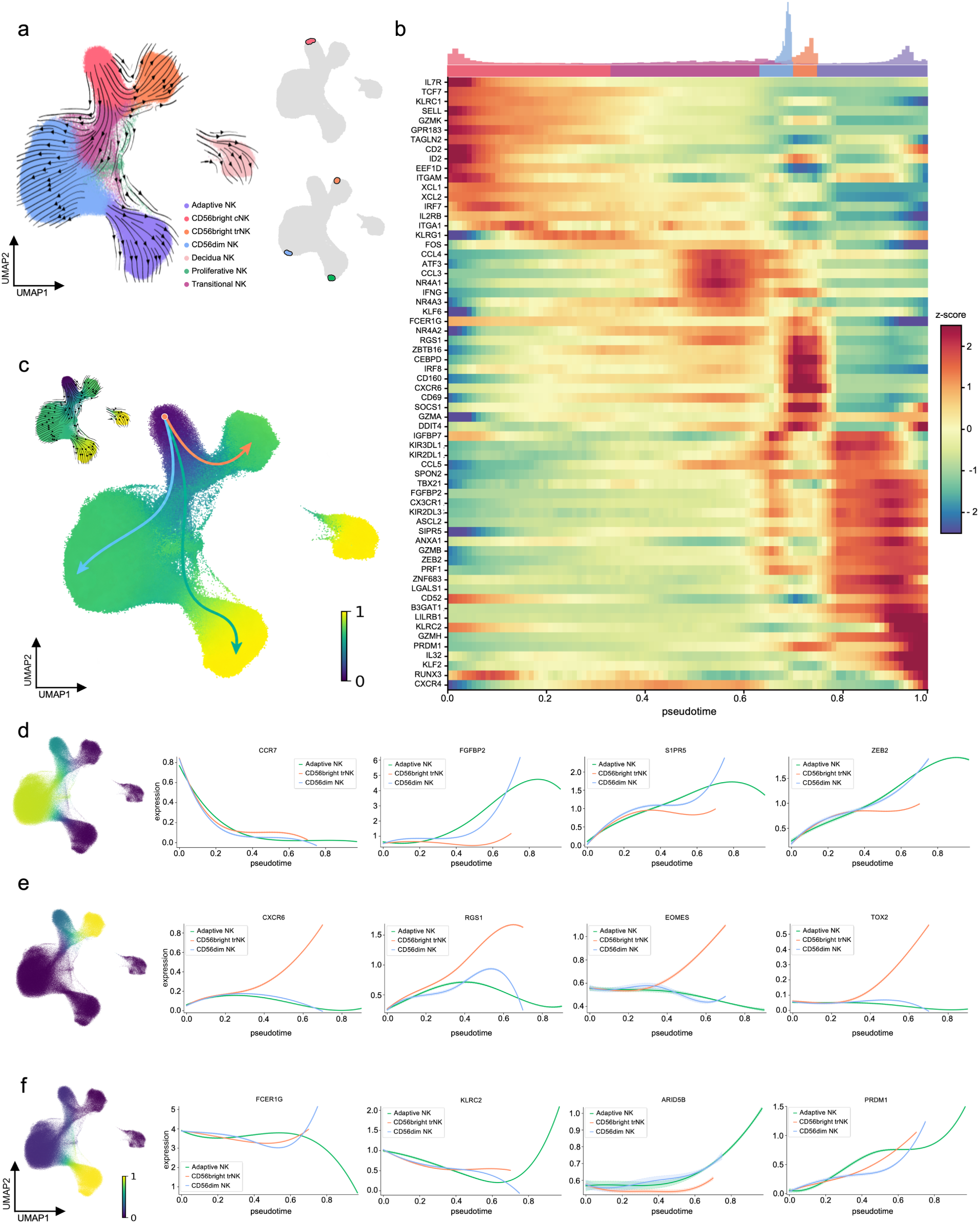
Differentiation trajectories embedded in the healthy NK-cell reference manifold. **a**, UMAP of the integrated HNKA colored by major NK-cell compartments. Initial and terminal macrostates are highlighted in the accompanying UMAPs. The stream arrows visualize the inferred flow of cell states. **b**, Heatmap of expression of NK lineage-associated genes along pseudotime. Density tracks above the heatmap show the distribution of the major NK-cell compartments. **c**, Inferred differentiation paths from CD56bright cNK cells projected on the reference embedding. **d–f**, Lineage-specific gene expression profiles along pseudotime separately for CD56dim NK (d), CD56bright trNK (e) and adaptive NK cells (f).

**Extended Data Fig. 3.**
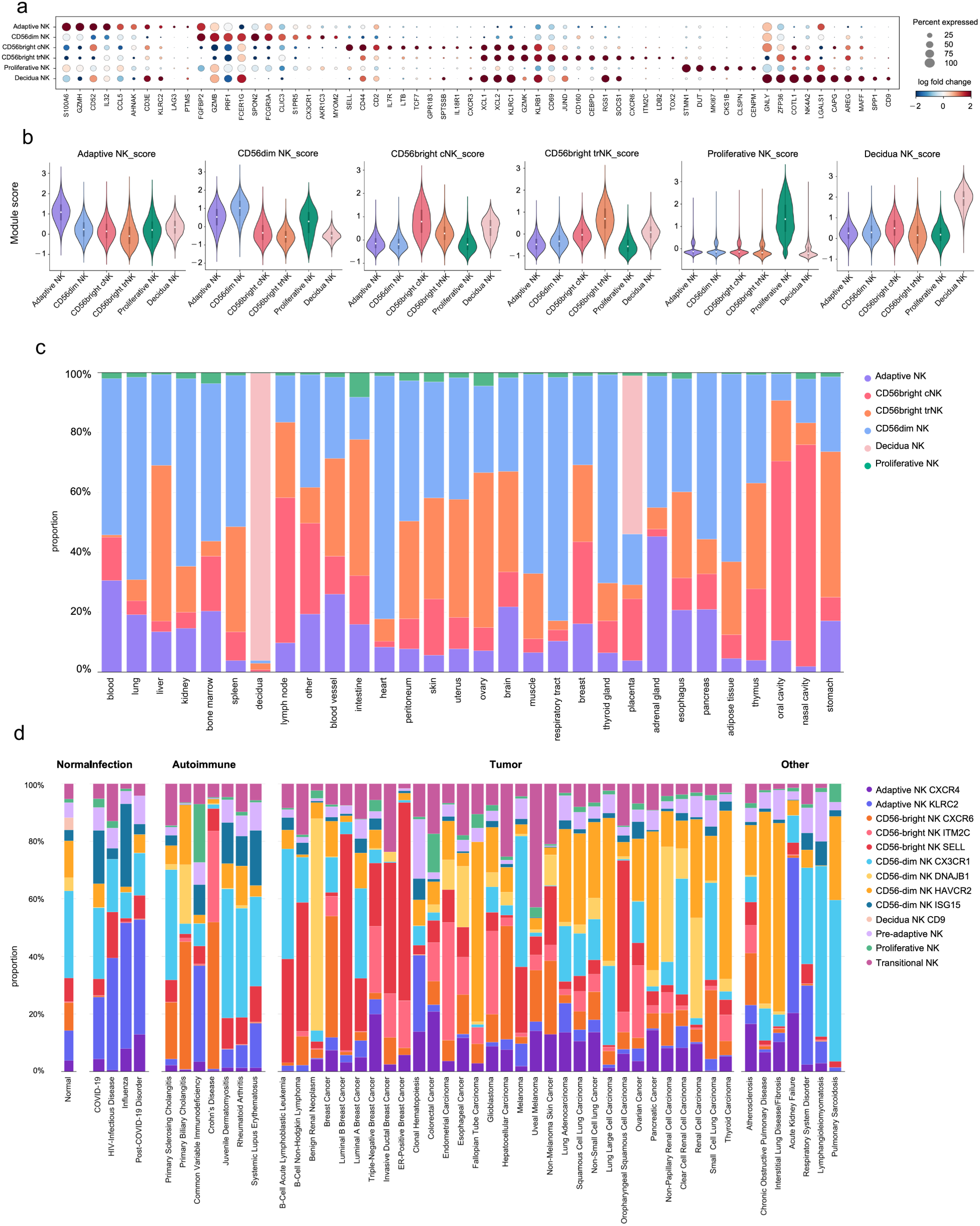
Marker specificity and compositional variation across tissues and diseases. **a**, Dot plot of representative marker genes across the major NK-cell compartments. **b**, Distributions of compartment-specific marker-module scores across the major NK-cell compartments. **c**, Relative abundance of the major NK-cell compartments. **d**, Relative abundance of the fine-grained NK-cell states across disease conditions.

**Extended Data Fig. 4.**
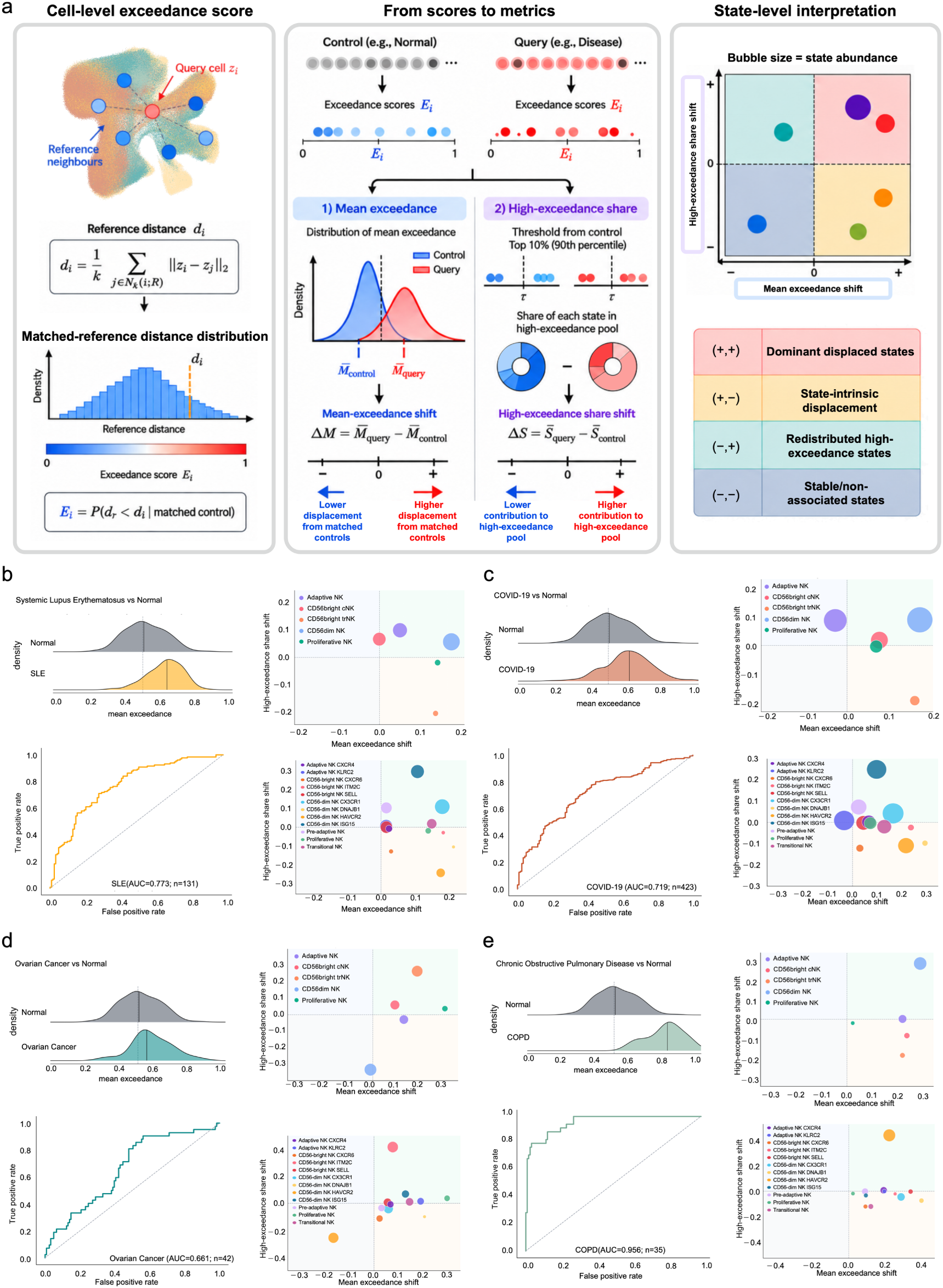
Exceedance framework and disease-level decomposition of NK-cell displacement. **a**, Schematic of the exceedance framework. Cell-level exceedance quantifies the distance of each query cell from tissue- and state-matched neighbors relative to the corresponding reference-distance distribution. Group-level differences are summarized by the mean-exceedance shift (ΔM) and the high-exceedance-share shift (ΔS), with high-exceedance cells defined relative to the 90th percentile of matched healthy controls. State abundance is represented by point size in the state-level decomposition. **b–e**, Exceedance in systemic lupus erythematosus (SLE; b), coronavirus disease 2019 (COVID-19; c), ovarian cancer (OV; d) and chronic obstructive pulmonary disease (COPD; e) relative to matched healthy samples. Density plots show donor-level mean exceedance, and receiver operating characteristic curves evaluate discrimination between disease and healthy donors; areas under the curve and numbers of disease donors are indicated. Bubble plots decompose the corresponding mean-exceedance and high-exceedance-share shifts across major NK-cell compartment (upper right) and fine-grained NK-cell states (lower right). Point area denotes relative state abundance and colors correspond to atlas annotations.

**Extended Data Fig. 5.**
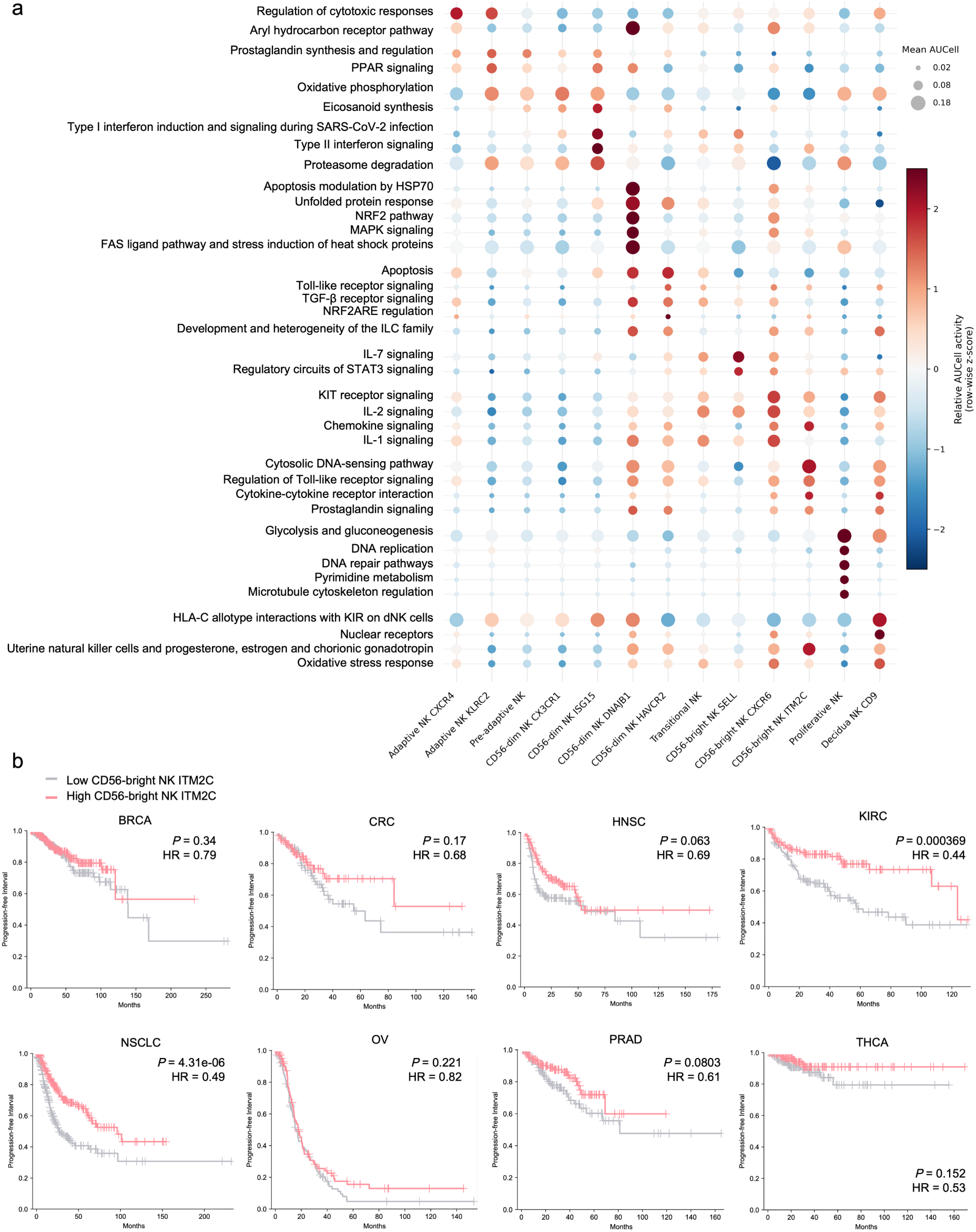
Functional programs of fine-grained NK-cell states and prognostic associations of CD56bright NK ITM2C. **a**, WikiPathways activity across fine-grained NK-cell states quantified by AUCell. Dot size denotes the mean AUCell score and color indicates row-wise z-score-normalized relative pathway activity across states. Selected pathways highlight state-associated cytotoxic, interferon-response, stress-response, cytokine-signaling, metabolic, proliferative and tissue-adaptation programs. **b**, Kaplan–Meier estimates of progression-free interval according to the CD56bright NK ITM2C signature in breast cancer (BRCA), colorectal cancer (CRC), head and neck squamous cell carcinoma (HNSC), kidney renal clear cell carcinoma (KIRC), non-small-cell lung cancer (NSCLC), ovarian cancer (OV), prostate adenocarcinoma (PRAD) and thyroid carcinoma (THCA). Tick marks indicate censored observations; P values and hazard ratios (HRs) are shown.

**Extended Data Fig. 6.**
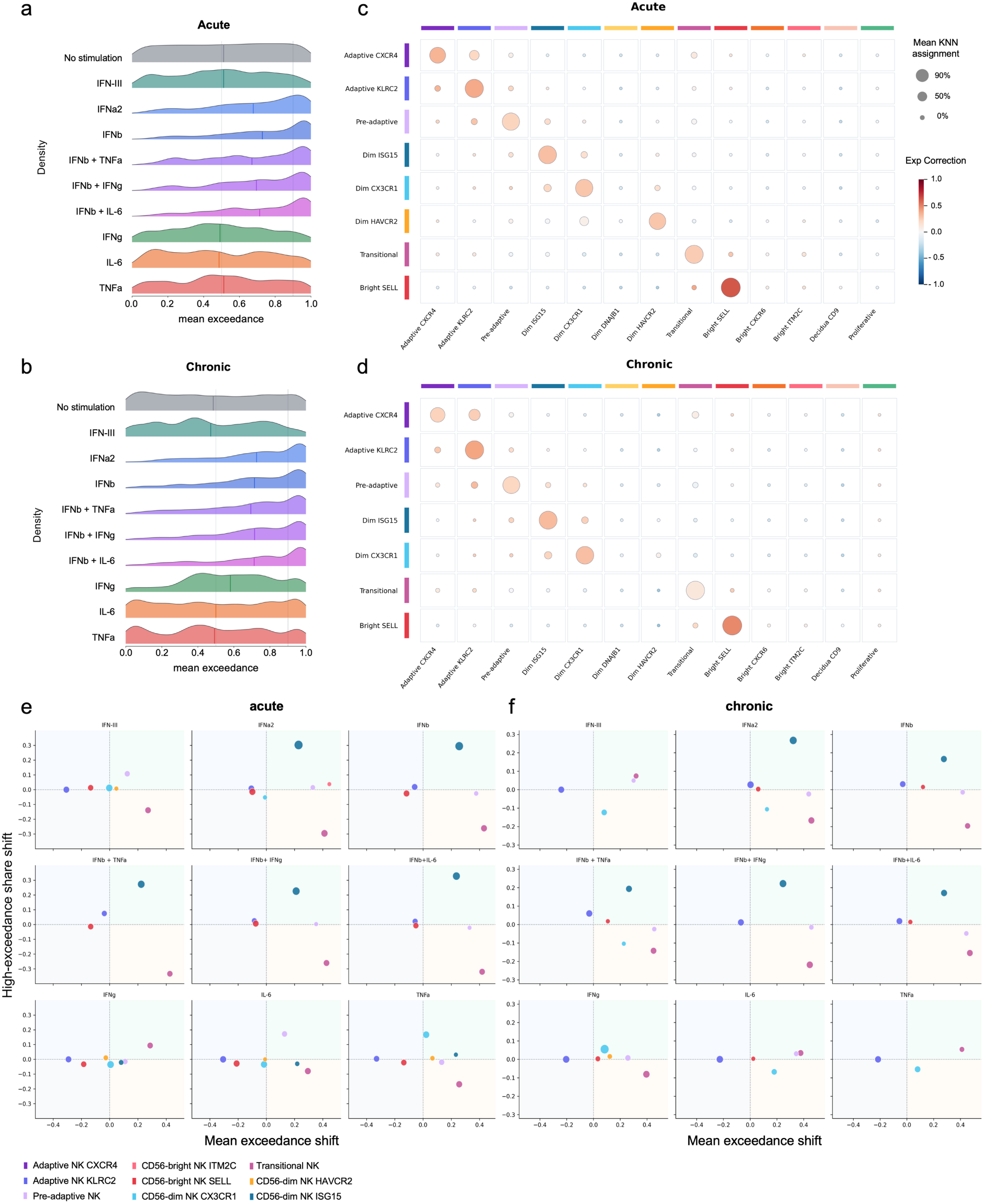
Atlas correspondence and reference-relative displacement of cytokine-perturbed NK cells. **a,b**, Distributions of cell-level exceedance following acute or chronic stimulation. Vertical lines indicate the threshold used to define high-exceedance cells. **c,d**, Correspondence between perturbation-mapped NK-cell states and atlas states after acute (c) or chronic (d) stimulation. Dot size denotes the donor/sample-unit-balanced soft-KNN assignment proportion, and color indicates expression-profile correlation calculated using atlas-centered, state-informative highly variable genes. **e,f**, Decomposition of exceedance after individual acute (e) or chronic (f) cytokine perturbations. Each point represents one NK-cell state; the x axis shows the shift in mean exceedance and the y axis shows the change in its share of high-exceedance cells. Point area denotes relative state abundance and colors correspond to the atlas annotations. KNN, k-nearest neighbor.

**Extended Data Fig. 7.**
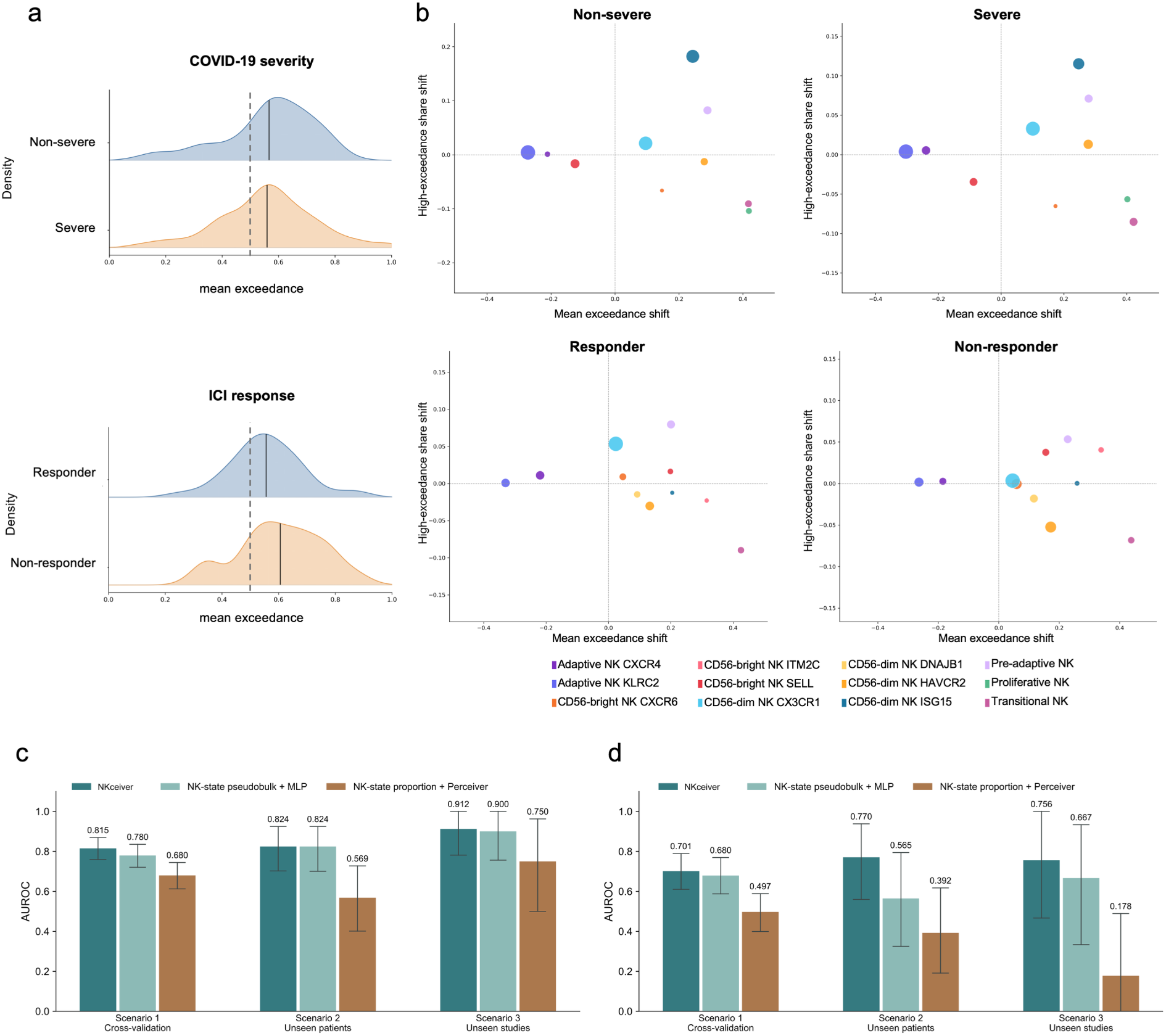
Decomposition of NK-cell displacement in COVID-19 severity and ICI response. **a,b**, Exceedance in COVID-19 severity and ICI response. Density plots show donor-level mean exceedance. Bubble plots decompose the corresponding mean-exceedance and high-exceedance-share shifts across NK-cell states. Point area denotes relative state abundance and colors correspond to atlas annotations. **c,d**, COVID-19 severity (c) and ICI response (d) performance across cross-validation, unseen patients and unseen studies.

